# Reference-based annotation of single cell transcriptomes identifies a profibrotic macrophage niche after tissue injury

**DOI:** 10.1101/284604

**Authors:** Dvir Aran, Agnieszka P. Looney, Leqian Liu, Valerie Fong, Austin Hsu, Paul J. Wolters, Adam Abate, Atul J. Butte, Mallar Bhattacharya

## Abstract

Myeloid cells localize to peripheral tissues in a wide range of pathologic contexts. However, appreciation of distinct myeloid subtypes has been limited by the signal averaging inherent to bulk sequencing approaches. Here we applied single-cell RNA sequencing (scRNA-seq) to map cellular heterogeneity in lung fibrosis induced by bleomycin injury in mice. We first developed a computational framework that enables unbiased, granular cell-type annotation of scRNA-seq. This approach identified a macrophage subpopulation that was specific to injured lung and notable for high expression of *Cx3cr1*+ and MHCII genes. We found that these macrophages, which bear a gene expression profile consistent with monocytic origin, progressively acquire alveolar macrophage identity and localize to sites of fibroblast accumulation. Probing their functional role, *in vitro* studies showed a trophic effect of these cells on fibroblast activation, and ablation of *Cx3cr1*-expressing cells suppressed fibrosis *in vivo.* We also found by gene set analysis and immunofluorescence that markers of these macrophages were upregulated in samples from patients with lung fibrosis compared with healthy controls. Taken together, our results uncover a specific pathologic subgroup of macrophages with markers that could enable their therapeutic targeting for fibrosis.

## Main

Fibrosing diseases are a major cause of mortality and, whether toxin-related, infectious, autoimmune, or idiopathic, can affect any organ^1^. Lung fibrosis is a particularly vexing clinical problem because of the lack of effective therapies and a poor understanding of its etiology. The most prevalent form, idiopathic pulmonary fibrosis, has a median survival of only 3 years, and approved treatments are limited^2^. With respect to pathogenesis, much attention has focused on fibroblast activation, given the central role of fibroblasts in the deposition of matrix proteins such as collagen^3-6^. However, the causes of fibroblast activation and proliferation are not fully understood.

The healthy adult lung contains both a self-renewing population of embryonically-derived macrophages known as alveolar macrophages and a separate population of interstitial macrophages present near the larger airways and in the lung interstitium^7,8^. In lung fibrosis models, macrophages expand in number and have been found to be profibrotic overall based on studies of broad-based ablation, such as with clodronate and pan-macrophage or developmental knockouts^9,10^. We used scRNA-seq to capture macrophage heterogeneity in the fibrotic phase of bleomycin injury and thus to enhance detection of functional subsets in fibrogenesis.

To facilitate analysis of mixed populations present in cell suspensions derived from whole lung, we developed SingleR (Single Cell Recognition), a reference-based computational tool that enables unbiased annotation of scRNA-seq. Whereas the prevalent approach in scRNA-seq has been to classify clusters of cells using marker genes, manual annotation suffers from subjectivity and low resolution. Furthermore, traditional markers have often been developed for flow cytometry based on their robust protein expression; however, the expression of these markers at the mRNA level may not support their use for transcriptomic approaches, particularly given the low detection and frequent dropout with scRNA-seq. SingleR overcomes these limitations by assigning cell-type identity to single cells by unbiased comparison to reference datasets of pure cell types sequenced by microarray or RNA-seq. The SingleR pipeline (Figure 1a and Supplementary Information) first correlates the single cell transcriptome against selected reference datasets. SingleR then iteratively fine-tunes the correlations by reanalyzing the top correlated cell types with a reduced set of genes that differentiate between them, allowing granular annotation of cell type.

**Figure 1.**
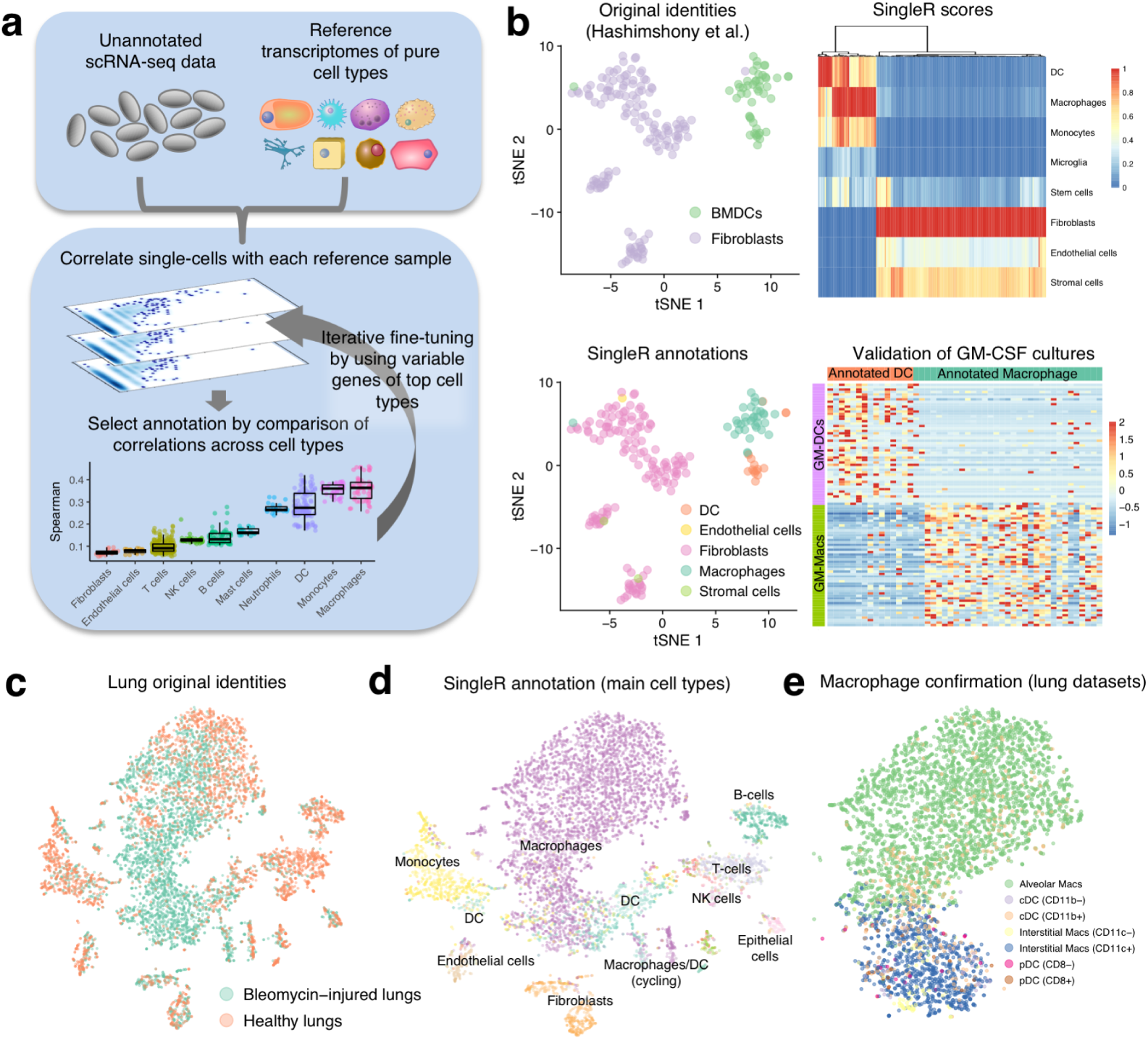
Annotation of scRNA-seq by reference datasets reveals disease-associated macrophages in lung fibrosis, **a**, Schematic of SingleR, a protocol for cell-type annotation by reference to transcriptomes of pure cell types (details in Supplementary Information), **b**, SingleR applied to a published scRNA-seq dataset of fibroblast and bone marrow-derived dendritic cells (BMDC)^11^. **Top left:** Original identities indicated on t-SNE plot. **Top right:** Heatmap of SingleR scores for top correlated cell types. **Bottom left:** SingleR annotation of cell identity indicated on t-SNE plot. **Bottom right:** Expression in single cells of genes derived by differential expression analysis of published microarray data from bone marrow-derived, GM-CSF-cultured macrophages (GM-Macs) and DCs (GM-DCs)^15^ confirms SingleR annotations, **c-e**, t-SNE plots of single-cell suspensions from mouse whole lung sequenced by Drop-seq and color-coded for experimental condition **(c)**, annotated by SingleR with the ImmGen reference database **(d)**, and annotated by SingleR with lung-specific myeloid datasets^24,25^ **(e).** Data shown pool replicates (n=3 mice for bleomycin, n=5 mice for control).

We first validated SingleR with several publicly available scRNA-seq datasets. For example, we studied scRNA-seq datasets of mouse bone marrow-derived dendritic cells (BMDC) and mouse fibroblasts published by Hashimshony et al.^11^ (Figure 1b). T-distributed stochastic neighbor embedding (tSNE) analysis of gene expression revealed separate clusters of fibroblasts and BMDCs conforming to their specified identities. We then applied SingleR using the ImmGen database of 830 microarray samples encompassing a broad range of pure mouse cell types^12^ as the reference library for unbiased annotation. Superimposing these annotations on the tSNE plot confirmed fibroblast identity but revealed, surprisingly, that 33 of the 48 BMDCs were macrophages. Of note, cells in the study were cultured with GM-CSF and sorted on CD11c for isolation; however, a recent study has shown that this procedure, which had been widely used in studies of DCs^13,14^, is not exclusive for DCs but also captures macrophages^15,16^. Comparative analysis using microarray data from this study confirmed segregation according to macrophage or DC lineage, in accordance with SingleR. These results show that SingleR enhances cell type annotation of scRNA-seq to a higher and more accurate resolution. Similar analyses for other datasets with comparisons of SingleR against alternative annotation methods^17,18^ as well as its application to >50 publicly available human and mouse scRNA-seq datasets are presented in Supplementary Information and in a dedicated web app http://comphealth.ucsf.edu/SingleR/.

We next applied SingleR to enhance analysis of transcriptomic change in lung fibrosis across multiple cell types by Drop-seq analysis of collagenase-digested whole mouse lungs, either at baseline or two weeks after bleomycin injury^19^. Altogether we sequenced 42,000 single-cells across five experiments, with 8,366 containing at least 500 non-zero genes, and used the Seurat package^20^ to perform tSNE for visualization of cellular clusters (Figure 1c, Supplementary Table 1, and Supplementary Figure 1). Using SingleR with the ImmGen database, we mapped the annotations to the tSNE plot (Figure 1d and Supplementary Figure 2). This analysis revealed a disproportionately large number of cells from bleomycin-injured lungs in a large central cluster (Figure 1c) that was annotated as macrophages (Figure 1d). Since macrophages and dendritic cells are known to have similar transcriptomic profiles under certain circumstances^21-23^, we repeated SingleR analysis using published RNA-seq datasets of mouse lung macrophages and dendritic cells^24,25^ for greater lung specificity. This analysis confirmed a predominance of lung macrophages as opposed to dendritic cells (Figure 1e and Supplementary Figure 3).

Two major annotations were resolved by this analysis: alveolar and interstitial macrophages. These annotations were strongest at the poles of our tSNE cluster, suggesting the possibility that there might be a continuum of gene expression between these macrophage cell types. Therefore, we quantified each cell by similarity analysis to the RNA-seq datasets of alveolar and interstitial macrophages^25^. This analysis revealed cells with intermediate gene expression between the two cell types (Figure 2a and Supplementary Figure 4). To better understand this intermediate cluster, we next used SingleR for unsupervised hierarchical subclustering of macrophages based on differential annotation using the full range of cell types within the ImmGen database and, consistent with our gradient analysis, identified three distinct groups: alveolar macrophages (C1), interstitial macrophages (C3), and an intermediate cluster of cells that were mapped by SingleR to both alveolar and non-alveolar macrophage reference datasets (C2; Figure 2b and Supplementary Figure 5).

**Figure 2.**
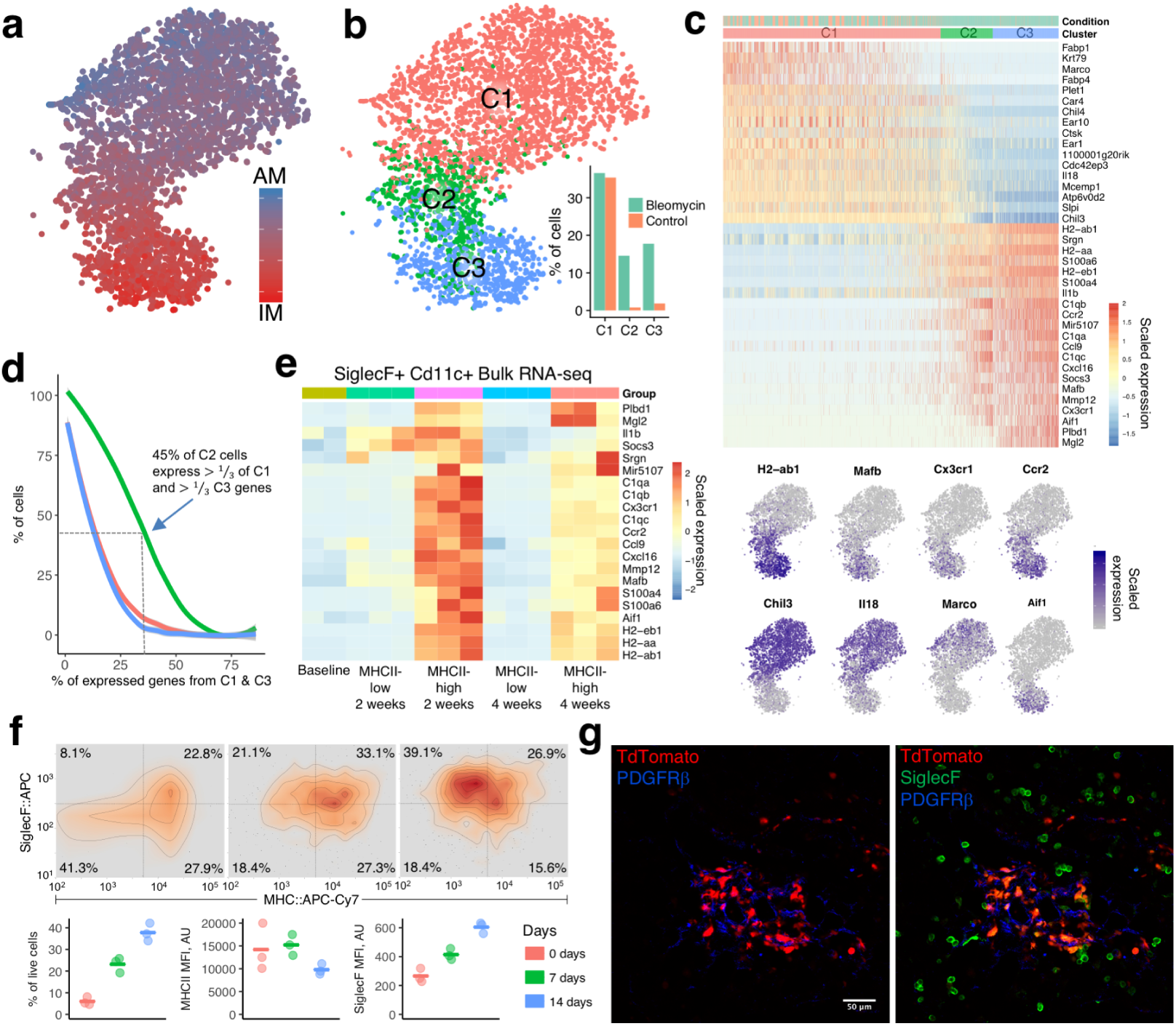
Macrophages expressing Cx3cr1 and MHCII localize to sites of fibroblast accumulation after lung injury, **a**, Quantification of similarity in gene expression of individual cells to bulk RNA-seq profiles of alveolar macrophages (AM) and CD11c+ interstitial macrophages (IM)^25^. b, Subclustering of macrophages based on SingleR annotation by reference to the ImmGen database, c, Heatmap of genes differentially expressed between C1 and C3 with examples superimposed on the tSNE plot, d, Percentage of cells in each cluster that express genes in c (differentially expressed between C1 and C3). 45% of C2 cells express at least a third of C1 genes and at least a third of C3 genes, e, Heatmap of the most differentially expressed genes from bulk RNA-seq of alveolar macrophages (SiglecF+CD11c+) sorted on MHCII-low at baseline and both MHC-low and MHCII-high at two time points after bleomycin (n=2 mice for baseline, n=3 mice at 2 weeks, n=3 mice at 4 weeks), f-g, Lineage tracing of lung cells in Cx3cr1-CreERT2 / Rosa26_l0xp_STOP_l0xp_-TdTomato mice with tamoxifen administration before and after injury followed by flow cytometry, with contour plots and box plots depicting values for TdTomato+ cells (f, n=3 mice for each group, mean is marked), and immunofluorescence (g, representative image is shown, n=3 mice).

Notably, both the intermediate subcluster C2 and C3 were highly enriched for cells from bleomycin-induced fibrosis. Differential gene expression between C1 and C3 and yielded 38 genes with log_2_ fold change > 1.5 (Figure 2c and Supplementary Table 2). Remarkably, a high percentage of cells from C2 had high expression of genes from both C1 and C3 (Figure 2d), suggesting that C2 represents a transitional state unique to the disease model and intermediate between C1 and C3. Moreover, *Cx3cr1, Ccr2, Mafb*, and MHCII genes were common to C2 and C3 and are consistent with macrophages of monocytic origin, as previously reported in bleomycin-induced lung fibrosis^10,26^. To confirm these findings for C2, we sorted alveolar macrophages (SiglecF+ CD11c+) after injury by MHCII-low and MHC-high expression to distinguish the C1 and C2 subclusters, respectively. MHCII-high cells compared with MHCII-low shared almost all the genes identified by scRNA-seq (Figure 2e and Supplementary Figure 6a).

We next undertook studies to probe the functional niche of these macrophages *in vivo*. Since *Cx3cr1* was expressed in both subclusters C2 and C3, we took advantage of the tamoxifen-inducible allele *Cx3cr1-CreERT2* ^27^ crossed with *Rosa26-loxp-STOP-loxp-TdTomato* for lineage tracing. Tamoxifen induction of healthy mice showed that a minority of lung cells, likely airway and interstitial macrophages, express *Cx3cr1* at baseline (Figure 2f and Supplementary Figure 6b). After injury, we observed an increase in *Cx3cr1*-lineage cells (CLCs). Furthermore, these cells progressively decreased expression of MHCII and increased expression of SiglecF, consistent with a transition of CLCs toward alveolar macrophage identity. Also consistent with a transition is the fact that, by bulk RNA-seq of alveolar macrophages (Figure 2e), the expression of C2 and C3 genes that were elevated at 2 weeks decreased toward baseline levels at 4 weeks after injury.

Surprisingly, we noted by confocal microscopy that CLCs were in direct contact with fibroblasts clustering at regions of injury and marked by expression of Pdgfrß (Figure 2g). Consistent with our scRNA-seq analysis, these CLCs also expressed the alveolar macrophage marker SiglecF, whereas non-lineage traced SiglecF+ macrophages (corresponding to the C1 subcluster) were absent from the fibrotic niche. Noting this localization of CLCs to sites of fibroblast accumulation, where scar forms in the model, we wondered whether these macrophages could provide trophic support for fibroblasts. This hypothesis is supported by recent *in vitro* evidence of the dependence of fibroblasts on macrophage-derived platelet derived growth factor (PDGF)^28^. Importantly, *Pdgfa* was upregulated in our bulk RNA-seq analysis of activated (MHCII-high) alveolar macrophages after injury (Figure 3a). To test whether CLCs support fibroblast migration or proliferation via Pdgfa, we collected conditioned medium from sorted alveolar macrophages and tested its effect on cultured fibroblasts in an *in vitro* gap closure assay. Conditioned media from MHCII-high alveolar macrophages sorted from injured lung significantly enhanced gap closure by 3T3 fibroblasts compared with MHCII-low conditioned media, an effect that was inhibited by antibody blockade of PDGF-AA (Figure 3b). To test the function of CLCs *in vivo*, we employed Diphtheria Toxin A-mediated ablation of cells expressing *Cx3cr1* during the fibrotic phase of bleomycin lung injury (days 8 through 21). Remarkably, CLC ablation decreased bleomycin-induced lung fibrosis by hydroxyproline assay (Figure 3c and Supplementary Figure 7). These results support a profibrotic effect of CLCs after injury and suggest that CLCs induce fibrosis via a trophic effect on fibroblasts.

**Figure 3.**
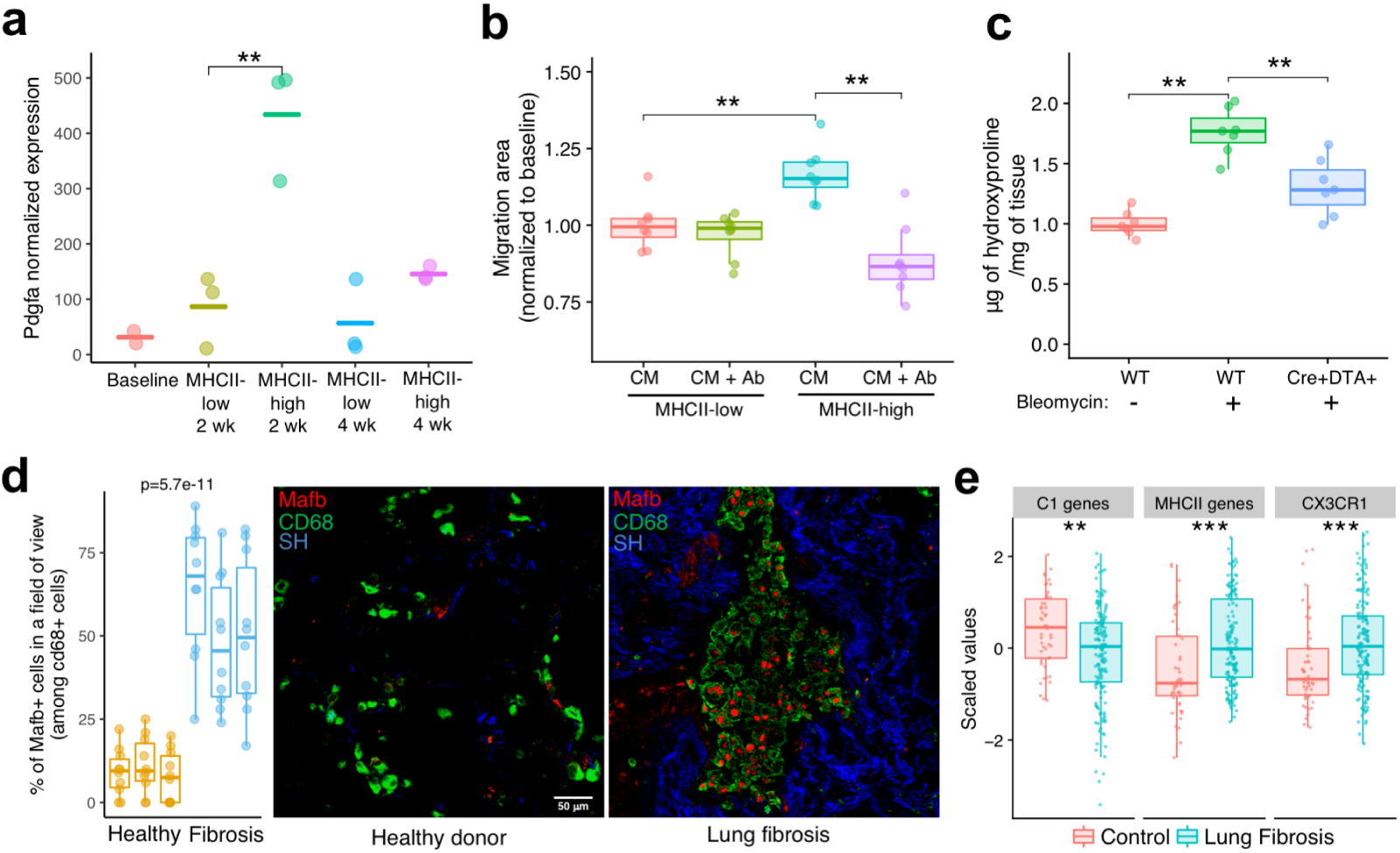
Cx3cri-derived macrophages are necessary for lung fibrosis and are upreguiated in human disease, **a**, *Pdgfa* expression by bulk RNAseq of SiglecF+CD11c+ lung macrophages sorted after bleomycin injury. Mean is marked. Wald test p-value is presented, **b**, 3T3 mouse fibroblast migration assay in presence of conditioned media (CM) from lung macrophages sorted by high and low MHCII expression 2 weeks after lung injury, with and without PDGF-AA blocking antibody (Ab; n=6 mice). Each data point corresponds to a separate assay, **c**, Lung collagen content measured by hydroxyproline assay 21 days after injury with and without macrophage ablation during the fibrotic period (day 8 onward) in *Cx3cr1-*CreERT2 x Rosa26_loxp_STOP_loxp_-Diptheria Toxin A mice (n=7 mice in each group), **d**, Representative immunofluorescence images of lung sections taken from human fibrotic and control lung with quantitation (10 fields of view per sample, n=3 fibrotic and 3 normal lung samples). SH=second harmonic imaging of collagen, **e**, Expression of human orthologues of the ImmGen alveolar (AM) signature, MHCII genes, and *CX3CR1* in bulk RNAseq samples from patient lung biopsy specimens^29^. Mean is shown in all boxplots, and Wilcoxon text p-values are presented. **p<0.01. ***p<0.001.

Finally, we tested the relevance of CLCs to human disease. Immunofluorescence of samples from patients with idiopathic lung fibrosis as opposed to healthy controls was notable for expression of MAFB, a marker unique to CLCs (Figure 2c), in CD68+ macrophages (Figure 3d). Furthermore, gene set analysis using human orthologues of murine genes identified in our study confirmed decreased alveolar macrophage gene expression and increased *CX3CR1* and MHC II genes in a published microarray dataset from patients with idiopathic pulmonary fibrosis compared to healthy controls (Figure 3e)^29^.

In summary, this study reveals a profibrotic macrophage subpopulation that localizes to sites of fibrotic scar with activating effects on the lung mesenchyme. Furthermore, the identification of specific markers for this group, including CX3CR1, could be used for therapeutic targeting in fibrosis. We also demonstrate the value of using reference-based annotation of single cell datasets in order to map the heterogeneity of canonical cell types such as macrophages, enhancing differentiation of transcriptomically unique subclusters within the data. Coupled with lineage tracing and functional studies, this approach should serve as a broadly applicable platform for study of specific cellular compartments in health and disease.

## Methods

### Mice

Ai14 (Rosa26-LSL-tdTomato), DTA (Gt(ROSA)26Sor^tm1(DTA)Lky^), CX3CR1CreERT (Cx3cr1^tm2.1(cre/ERT2)Jung^), all on C57BL/6 background, wild type C57BL/6, and wild type 129S1 mice were obtained from the Jackson Laboratory. Strain-appropriate controls were used in all experiments, and genotyping of the mice was performed by PCR. CX3CR1-CreERT2 mice were administered 2 mg tamoxifen (Sigma) via IP injection every other day for Cre induction. Mice were maintained in specific-pathogen-free conditions in the Animal Barrier Facility of the University of California, San Francisco. All animal experiments were approved by the Institutional Animal Care and Use Committee of the University of California, San Francisco.

### Microfabrication of Devices

The PDMS Drop-seq droplet microfluidic device is fabricated with standard soft lithography techniques. Photoresist masters are created by spinning a layer of photoresist SU-8 (Microchem) onto a 3 inch silicon wafer (University Wafer), then baking at 95°C for 20 minutes. Then, the photoresist is subjected to 3 minutes ultraviolet exposure over Drop-seq^19^ photolithography masks (CAD/Art Services) printed at 12,000 DPI. After ultraviolet exposure, the wafers are baked at 95°C for 10 min, developed in fresh propylene glycol monomethyl ether acetate (Sigma Aldrich), and rinsed with fresh propylene glycol monomethyl ether acetate, and baked at 95°C for 1 minute to remove solvent. The microfluidic devices are fabricated by curing poly(dimethylsiloxane) (10:1 polymer-to-crosslinker ratio) over the photoresist master^30^. The devices are cured in an 65°C oven for 2 hours and extracted with a scalpel, and inlet ports are added using a 0.75 mm biopsy core (World Precision Instruments). The device is bonded to a glass slide using O_2_ plasma treatment, and channels are treated with Aquapel (PPG Industries) to render them hydrophobic. Finally, the devices are baked at 65°C for 20 min to dry the Aquapel before they are ready for use.

### Drop-seq and data analysis

For scRNA-seq analysis of lung fibrosis, bleomycin (3 U/kg; Hospira) or water was instilled intratracheally to anesthetized male 129S1 mice, age 10-12 weeks. After 2 weeks mice were sacrificed, and lungs were perfused with PBS and dissociated to a single cell solution in RPMI containing 0.13 U of liberase TM (Roche) and dissociated using gentleMACS (Miltenyi Biotec). Cells were passed through a 70 um and 40 um strainers. The single cell RNA-seq experiment is performed based on the Drop-seq protocol^19^. Briefly, the barcoded Drop-seq beads (ChemGenes corporation, MACOSKO-2011-10) and single cell suspension from dissociated mouse lung are re-suspended to 100 beads/µl in PBS-BSA buffer and 120 cells/µl in Drop-seq lysis buffer (with additional 1 M NaCl added), respectively. Monodisperse droplets ∼1 nL in size were generated using the fabricated Drop-seq device. We used HFE7500 with 2% w/v ionic krytox as oil phase. The collected droplets were broken with perfluorooctanol (Sigma) in 30 ml of 6× SSC buffer. The beads were then washed and re-suspended in a reverse transcriptase mix for reverse transcription and the template switch reaction. Exonuclease I is used to remove unextended primers following the RT reaction. The beads were then washed, counted, aliquoted into PCR tubes as 6000 beads per PCR reaction, and PCR-amplified for 17 cycles: 95°C for 3 min; then 4 cycles of: 98°C for 20 sec, 65°C for 45 sec, 72°C for 3 min; then 13 cycles of: 98°C for 20 sec, 67°C for 20 sec, 72°C for 3 min; and finally, 72°C for 5 min. The PCR reactions from one Drop-seq run were pooled and purified with AMPure XP beads; the amplified cDNA was quantified with Qubit dsDNA high sensitivity assay and checked on a BioAnalyzer High Sensitivity Chip (Agilent). The cDNA was fragmented and amplified for sequencing with the Nextera XT DNA sample prep kit (Illumina) using a primer that enabled the specific amplification of only the 3’ ends and Illumina index primers N70X. The libraries were purified with Ampure beads, quantified with Quabit dsDNA high sensitivity assay, checked on a BioAnalyzer High Sensitivity Chip, and then sequenced on Illumina Miseq or Hiseq2000.

Paired-end sequence reads were processed mostly as previously described^19^. Briefly, cell barcode and UMI are extracted from Read 1. Read 2 was aligned to the mouse mm10 genome (UCSC) using Bowtie^31^. Reads mapping to exonic regions of genes of mouse mm10 genome were recorded. The number of transcripts for a given gene within a cell barcode was determined by counting unique UMIs and were compiled into a digital gene expression (DGE) matrix.

### Public gene expression data

Immunological Genome Project (ImmGen) raw expression data of phase 1 and 2 were downloaded as CEL files from GEO (GSE15907 and GSE37448), processed and normalized using the Robust Multi-array Average (RMA) procedure on probe-level data using Matlab functions. The analysis was performed using custom CDF file obtained from Brainarray^32^. Raw counts from of the lung reference dataset were obtained from Gibbins et al.^25^ (GSE94135) and Altboum et al.^24^ (GSE49932). Counts were normalized and converted to TPM using R functions. Gene expression profile of GM-CSF derived bone marrow dendritic cell subsets (available at GSE62361)^15^ was analyzed using the GEO2R tool for differentially expressed genes. Top 50 upregulated by fold-change of GM-DCs genes and top 50 GM-Macs were used in Figure 2b.

### Cell Type Annotation of Single Cells

Feature counts from 10 batches were combined (42,000 cells) and analyzed with the Seurat toolkit v2.2^19^. Cells with less than 500 were filtered out; genes expressed in only 1 cell were omitted. Altogether, the filtered data contained 8,366 cells and 13,861 genes (Table S1). Expression data was log-normalized, and data were scaled to regress out differences in number of detected molecules. Variable genes were identified using the FindVariableGenes function with a x.low.cutoff = 0.0125, x.high.cutoff = 3 and y.cutoff = 0.5. t-SNE plot was computed using the top 10 principal components. SingleR was run in ‘DE’ mode, choosing reference variable genes based on pairwise cell type differences in expression levels. For each single cell transcriptome, Spearman coefficients for correlations with multiple samples of the same cell type in the reference dataset were computed and separately binned by cell type. Annotation scores of the single cell for each cell type were the recorded as the 75^th^ percentile Spearman coefficient value for each cell type.

For each single-cell, top associated cell types (with <0.05 difference from maximal score) were then reanalyzed using only variable genes among those cell types. In each round, the cell type with the lowest correlation or cell types with >0.05 difference from maximal score were dropped. This process was iterated until a single cell type remained, which was rendered cell type annotation. More details about the method can be found in Supplementary Information.

Clustering in Figure 2b was performed using Ward’s hierarchical agglomerative clustering method on the SingleR scores across all cell types. Deconvolution analysis in Figure 2a was performed using the DeconRNAseq package^33^ using the average expression of AM samples and average expression of IM3 samples from GSE94135. Differential expression analysis (Figure 2c) was performed using the Seurat package v2.2 with default parameters.

### Single cell dissociation for FACS

Lungs were perfused with PBS and dissociated to a single cell solution in 10mM HEPES RPMI containing 0.2% collagenase (Wako Pure Chemical Industries), 0.1 mg/mL Dispase II (Roche), and 2000 U/mL DNase I (Merck). Cells were passed through a 70 um strainer and stained at 4 °C for 30 minutes with following antibodies (1:100): SiglecF-APC (clone 1 RNM44N, eBioscience), MHCII-APC-Cy7 (clone M5/114.15.2, eBioscience), CD11c-PE (clone HL3, BD Biosciences). Cells were sorted either with BD FACSAria2 or SONY SH800 FACS. Data for analytical flow was analyzed using FlowJo software.

### Bulk RNA-seq

For bulk RNA-seq analysis of lung fibrosis, bleomycin (3 U/kg; Hospira) or water was instilled intratracheally to anesthetized male 129S1 mice, age 10-12 weeks. Cells gated on SiglecF+CD11c+ and sorted on MHCII-high or -low (Supplemental Figure S7) were collected at 2 weeks and 4 weeks after injury, or at baseline for the MHC-low compartment. RNA was collected (RNeasy Micro, Qiagen) from these samples, and total RNA quality was assessed by spectrophotometer (NanoDrop, Thermo Fisher Scientific Inc., Waltham, MA) and the Agilent 2100 Bioanalyzer (Agilent Technologies, Palo Alto, CA). Intact mRNA was isolated using the Dynabead mRNA Purification Kit for total RNA, according to manufacturer’s protocol (Thermo Fisher Scientific, Waltham, MA). Amplified cDNA was prepared using the NuGen Ovation RNA-Seq system V2 kit, according to the manufacturer’s protocol (NuGen Technologies, Inc., San Carlos, CA), and sequencing libraries were generated using the Nextera XT library preparation kit with multiplexing primers according to manufacturer’s protocol (Illumina, San Diego, CA). Library fragment size distributions were assessed using the Bioanalyzer 2100 and the DNA high-sensitivity chip (Agilent Technologies, Santa Clara, CA). Library sequence quality was assessed by sequencing single-end 50 base pair reads using the Illumina MiSeq platform and were pooled for high-throughput sequencing on the Illumina HiSeq 4000 by using equal numbers of uniquely mapped protein coding reads. RNA sequencing was performed on HiSeq 4000 machines (Illumina, San Diego, CA), followed by de-multiplexing of raw sequencing results, trimming of adapter sequences, and alignment to reference genome using STAR software^34^. DESeq2 was used to normalize by size factor (reads per sample) as well as by library complexity and then applied Wald test to determine significance of differential expression.

### Immunostaining and Fluorescence Microscopy

For immunostaining mouse lungs were perfused with PBS, inflated with 50% OCT, 10% sucrose in PBS and fixed overnight at 4 C in 4% PFA. For human samples OCT embedded tissues were cut to 15um sections and sections were fixed with 4% PFA for 15 minutes at RT. OCT embedded tissues were cut to 15um sections, blocked with 3% donkey serum, 1% BSA, 0.3% TritonX and then stained with following antibodies (1:100): SiglecF (clone 1RNM44N, eBioscience or AF1706, R&D Systems), Mertk (AF591, R&D Systems), PDGFRb (clone APB5, eBioscience), CD68 (KP1, Abcam or PA1518, Boster), MafB (HPA005653, Sigma). Secondary antibodies conjugated to Alexa fluorophores (Invitrogen and Abcam) were used at concentration 1:200. Stained sections and 2-photon images of second harmonic signal were visualized at Zeiss LSM 780 NLO microscope. Image processing was done using ImageJ.

### Fibroblast Migration Assay

3T3 mouse fibroblasts were seeded at 100% confluency into a 96-well plate central cell-free detection zone in the center of each well (Platypus technologies). 100 µl of conditioned media of macrophage cultures (48h culture in RPMI containing 10% FBS) were added per well. PDGF-AA antibody (07-1436, Millipore) was added at concentration 0.05 µg/ml. Cells were imaged via bright field microscopy at time points: 0h and 24h. Image processing and migration quantification was done using ImageJ.

### Measurement of hydroxyproline

Lung hydroxyproline content was measured as previously described^6^ three weeks after intratracheal instillation of bleomycin to sex-matched, male and female, 8-12-week-old C57BL/6J mice. Briefly, mouse lungs were incubated in 12 N HCl at 110°C for 18 h. Aliquots of the samples reconstituted in distilled water were added to 1.4% chloramine-T in 10% isopropanol and 0.5 M sodium acetate. Erlich’s solution was added, and the samples were incubated at 60°C for 10 min. Absorbance at 562 nm was measured and adjusted according to standard curves.

### Human Lung Tissues

Written, informed, consent was obtained from all subjects and the study approved by the UCSF institutional review board. Pulmonary fibrosis lung tissues were obtained at the time of lung transplantation from patients with a pathologic diagnosis of usual interstitial pneumonia and a consensus clinical diagnosis of idiopathic pulmonary fibrosis (IPF) according to available guidelines^35^. Age-similar, non-diseased, normal lung tissues were procured from lungs not used by the Northern California Transplant Donor Network. Our studies indicate that these lungs are physiologically and pathologically normal^36^. Lung fragments were inflated with 30% OCT suspended in PBS, embedded in OCT compound and frozen immediately after isolation in dry ice before storing at −80 °C until use.

### Analysis of published microarray datasets

Yang et al.^29^ performed gene expression profiling for 167 lung tissues from fibrosing idiopathic interstitial pneumonia subjects and 50 healthy controls. We downloaded raw CEL files GEO (Accession GSE32537) and processed using custom CDFs from BrainArray. Normalization was performed using the Robust Multi-array Average (RMA) procedure on Affymetrix microarray data. Single-sample gene set enrichment analysis (ssGSEA)^37^ was performed using human orthologues of C1 genes (Extended Data: Table S3) and 15 human MHCII genes (HLA-DMA, HLA-DMB, HLA-DOA, HLA-DOB, HLA-DPA1, HLA-DPB1, HLA-DPB2, HLA-DQA1, HLA-DQA2, HLA-DQB1, HLA-DQB2, HLA-DRA, HLA-DRB1, HLA-DRB5, HLA-DRB6).

## Statistical analysis

All significance tests in this paper, unless otherwise stated, were assessed using the two-sided Wilcoxon rank-sum test.

## Data Availability

Raw reads and processed data of single-cell RNA-seq experiments were deposited to GEO, accession GSE111664 (https://www.ncbi.nlm.nih.gov/geo/query/acc.cgi?acc=GSE111664). Bulk RNA-seq expression profiles of SiglecF+Cd11c+ lung macrophages were deposited to GEO, accession GSE111690 (https://www.ncbi.nlm.nih.gov/geo/query/acc.cgi?acc=GSE111690). The SingleR package and R scripts for producing the figures are available at https://github.com/dviraran/SingleR (under the GPL 3.0 license).

## Acknowledgements

This work was supported by a UCSF Marcus Award to M.B. and A.A., a National Institutes of Health grant (HL131560) to M.B, a Gruss Lipper Postdoctoral Fellowship to D.A., and the National Institute of Allergy and Infectious Diseases (Bioinformatics Support Contract HHSN272201200028C) to A.J.B. The content is solely the responsibility of the authors and does not necessarily represent the official views of the National Institutes of Health. We thank D. Erle, A. Barczak, W. Eckalbar, and M Adkisson of the UCSF Functional Genomics Core Facility and D. Sheppard for his insightful comments on the manuscript.

## Author Contributions

D.A. developed the cell type annotation tool presented and performed computational analysis of single cell data under the guidance of A.B; A.L. performed in vivo and in vitro experiments on macrophage lineage and function under the guidance of M.B; L.L. performed microfluidic capture of single cell transcriptomes, library preparation, and sequencing under the guidance of A.A; V.F. and A.H prepared breeding and experimental stocks of genetically modified mice and performed lung injury models under the guidance of M.B.; P.J.W. contributed acquisition, storage, and processing of human samples; M.B. conceived of the work, supervised experimental planning and execution, and wrote the manuscript with input from D.A., A.L., and L.L.

