## Supplementary Materials for "Reference-based annotation of single cell transcriptomes identifies a profibrotic macrophage niche after tissue injury"

### Extended Data: Figures and Tables

##### Supplementary Figure 1

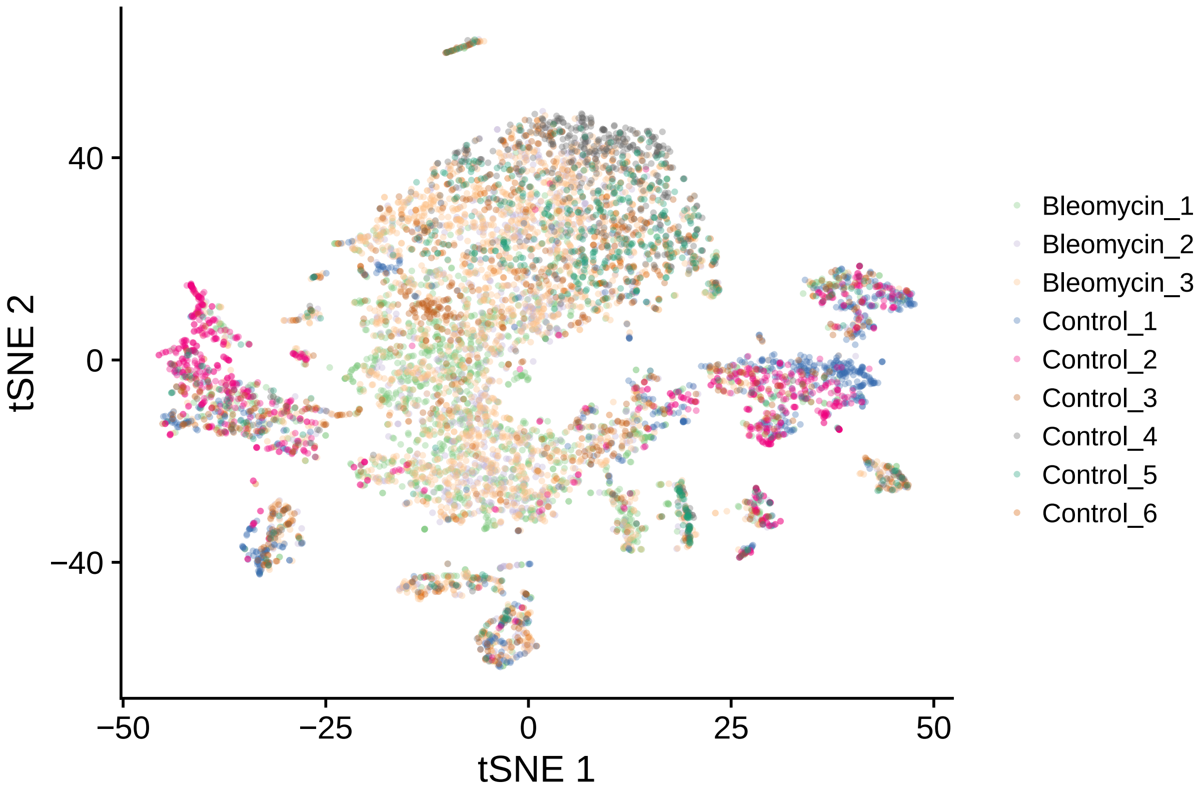

**Figure S1. 8,366 single-cells colored by batches.** This t-SNE plot accompanies Figure 1c. The colors are coded by batch. Within experimental condition (bleomycin or control) the batches intersect, suggesting that our data does not suffer from a strong batch effect.

#### Supplementary Figure 2

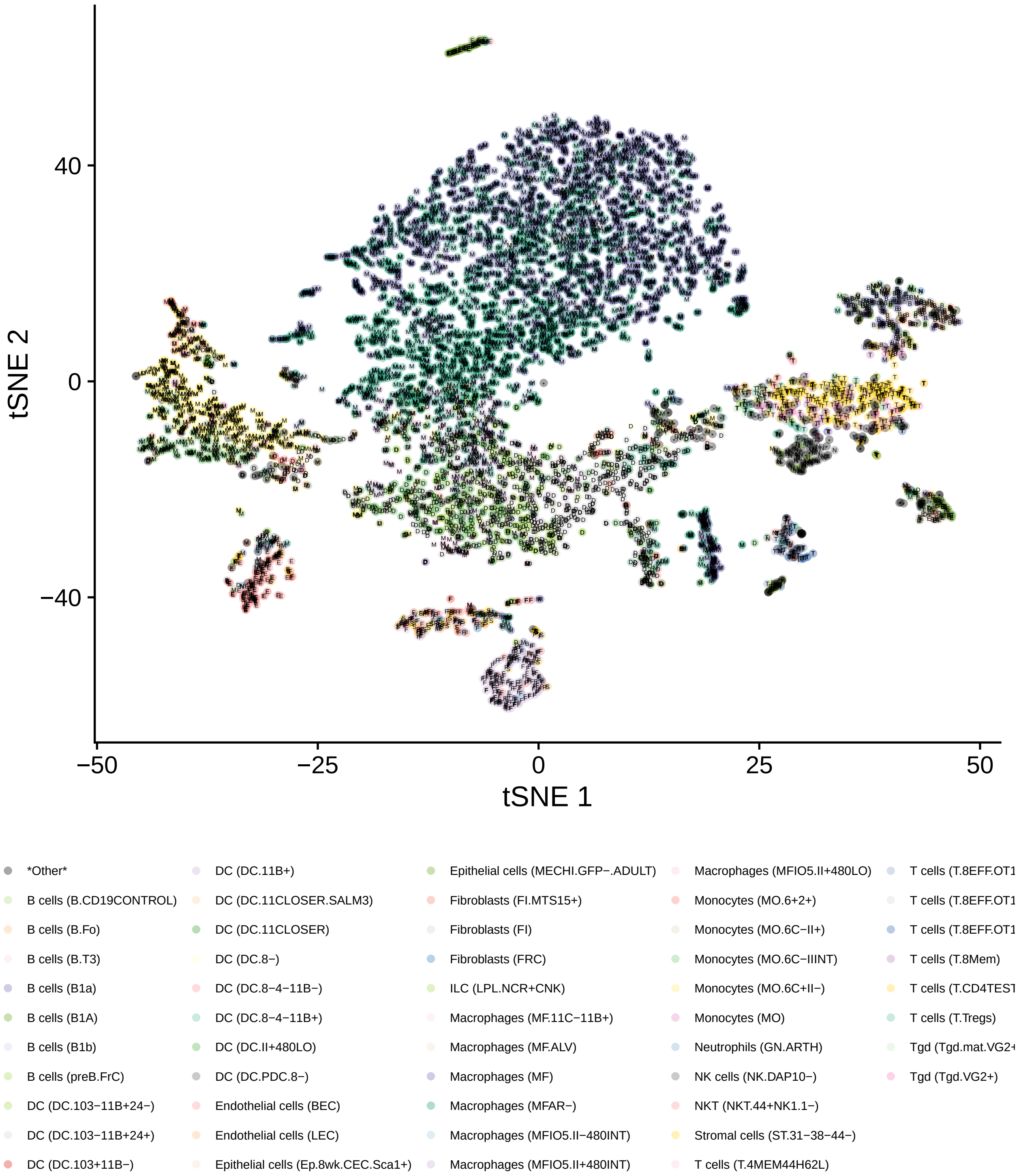

**Figure S2. SingleR detailed annotations.** This heatmap accompanies Figure 1d. Here we present all SingleR annotations (not just main types). Interestingly, SingleR annotated the cells in the middle of the macrophage cluster as alveolar macrophages extracted and probed in bulk by microarray from *pparg-/-* mouse, as opposed to the top cells, which were annotated as WT alveolar macrophages. To visualize the data, annotations with less than 15 cells were marked as *Other* and colored in black.

#### Supplementary Figure 3

**a**

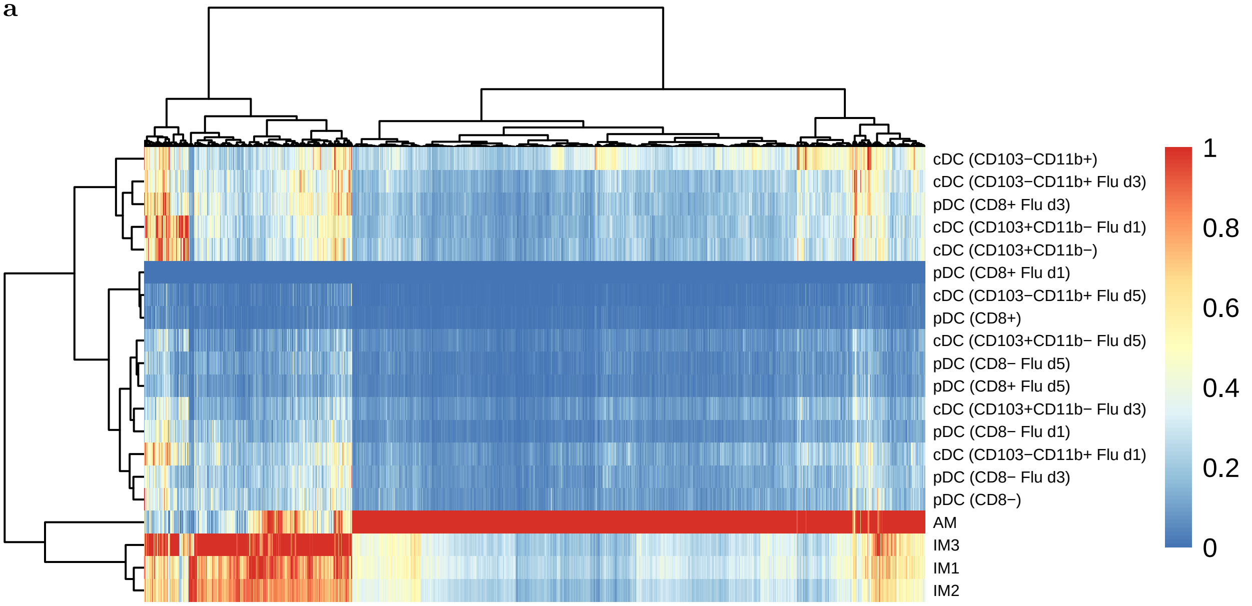

**b**

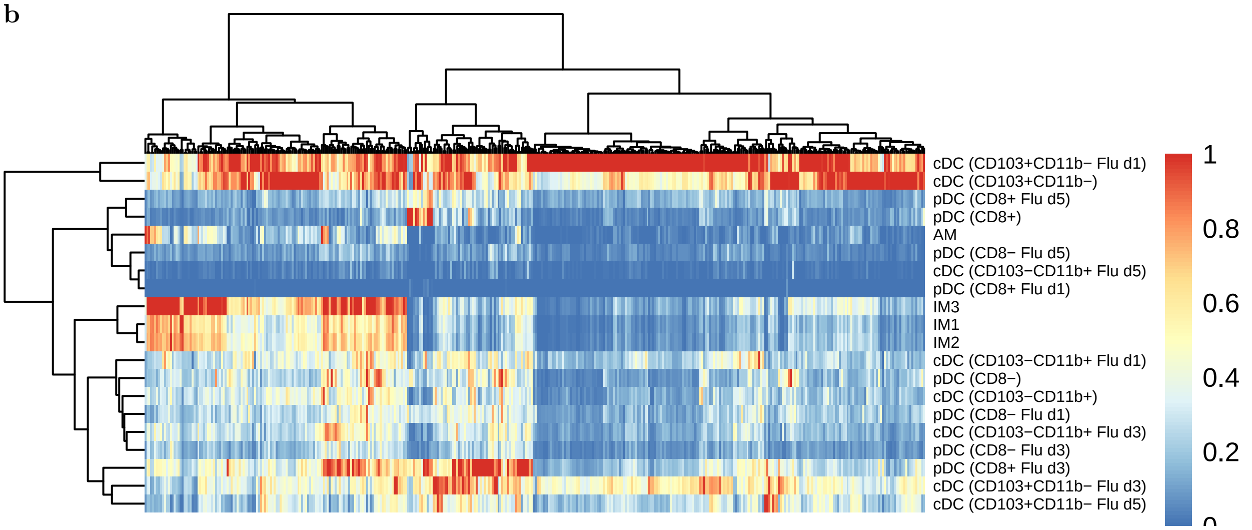

**Figure S3. Heatmap of SingleR scores using lung reference datasets. a.** This heatmap accompanies Figure 1e. We use RNA-seq datasets as reference - lung macrophages from Gibbings et al. (2017) (downloaded from GEO accession number GSE94135) and lung dendritic cells from Altboum et al. (2014) (downloaded from GEO accession number GSE49932). The two datasets were combined for a lung specific reference. We see that the majority of cells were clearly annotated to alveolar macrophages (AM), whereas the remaining were most correlated with IM3, which is CD11c+ interstitial macrophages, and not dendritic cells. Interestingly, the alveolar cluster was split to two clusters, as we show in Figure 2b. **b.** Cells from the dendritic cell cluster (see Figure 1d) are annotated as dendritic cells, showing that the annotations in (a) are not an artifact of the reference datasets.

**Supplementary Figure 4**

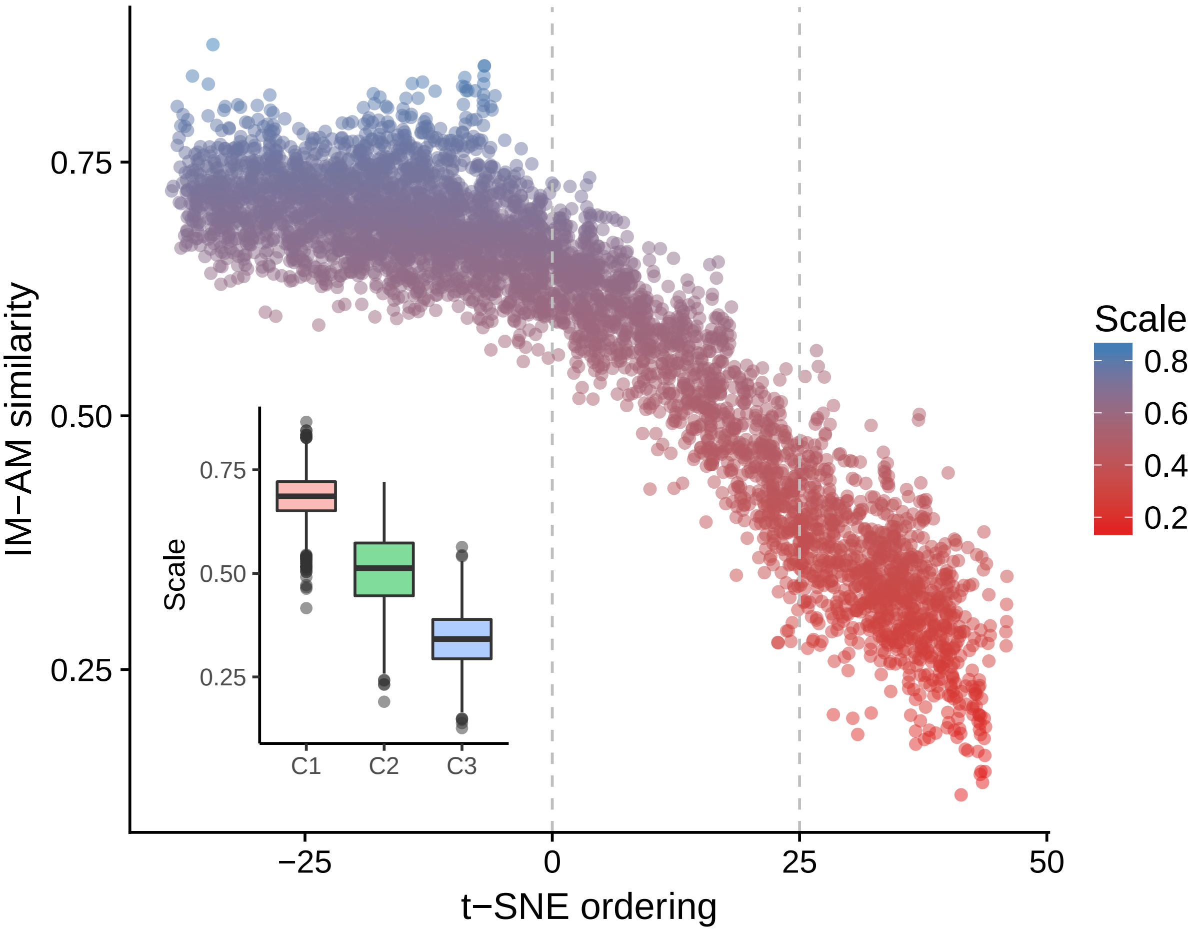

**Figure S4. Similarity analysis to alveolar and interstitial macrophages.** This plot accompanies Figure 2a. Similarity to alveolar macrophages (AM) and interstitial macrophages (IM) was quantified using a deconvolution approach. Using the DeconRNAseq package (Gong and Szustakowski 2013), we measured in each single cell the enrichment of genes along a continuum between AM and IM gene sets derived from lung macrophage reference data (GSE94135 used for Figure 1e). Here, 0 represents similarity to IM and 1 similarity to AM, and a value of 0.5 suggests an intermediate state. In this plot we present the value of IM-AM similarity for each cell as a function of a one-dimension t-SNE plot (the horizontal axis here is the first PCA computed on tSNE1 and tSNE2 from the original tSNE plot). The insert shows a boxplot of the AM-IM scale by the clusters as defined in Figure 2b. The dashed lines roughly distinguish the cells of each cluster. We observe a linear change from high similarity to AM towards high similarity to IM, suggesting a gradient of differentiation along AM to IM (or vice versa).

#### Supplementary Figure 5

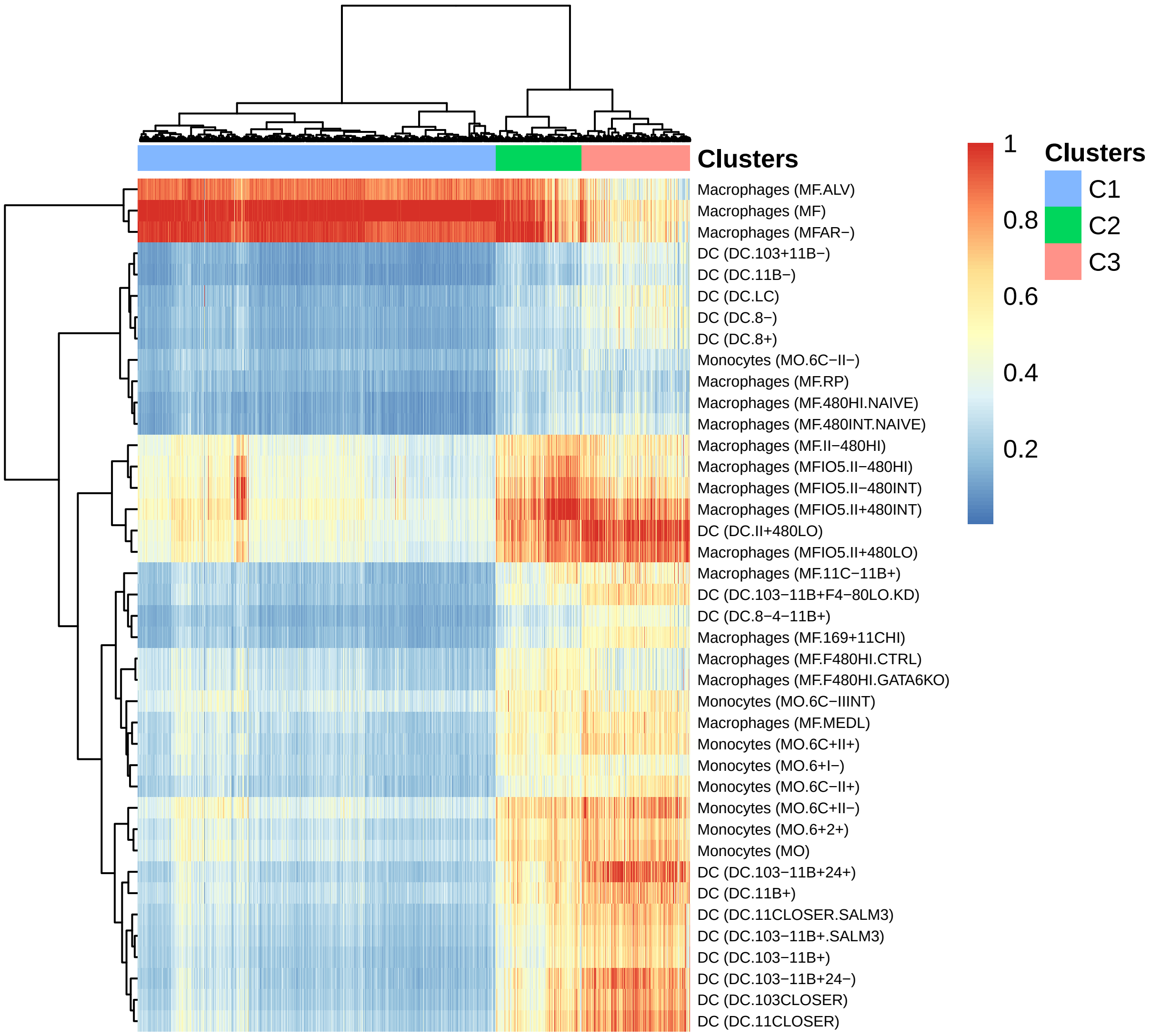

**Figure S5. Clustering lung macrophages using SingleR scores.** This heatmap accompanies Figure 2b. Using the ImmGen reference datasets we cluster the cells with correlation scores produced by SingleR, thus generating a heatmap based on differential annotation. Cluster 1 is highly associated exclusively with ImmGen alveolar macrophages. Cluster 3 is mainly associated with non-alveolar myeloid transcriptomes (which were shown in lung reference datasets to be interstitial macrophages in Figure 1e and Supplementary Figure 3). Cluster 2 is associated with both alveolar and non-alveolar macrophage myeloid cells.

#### Supplementary Figure 6

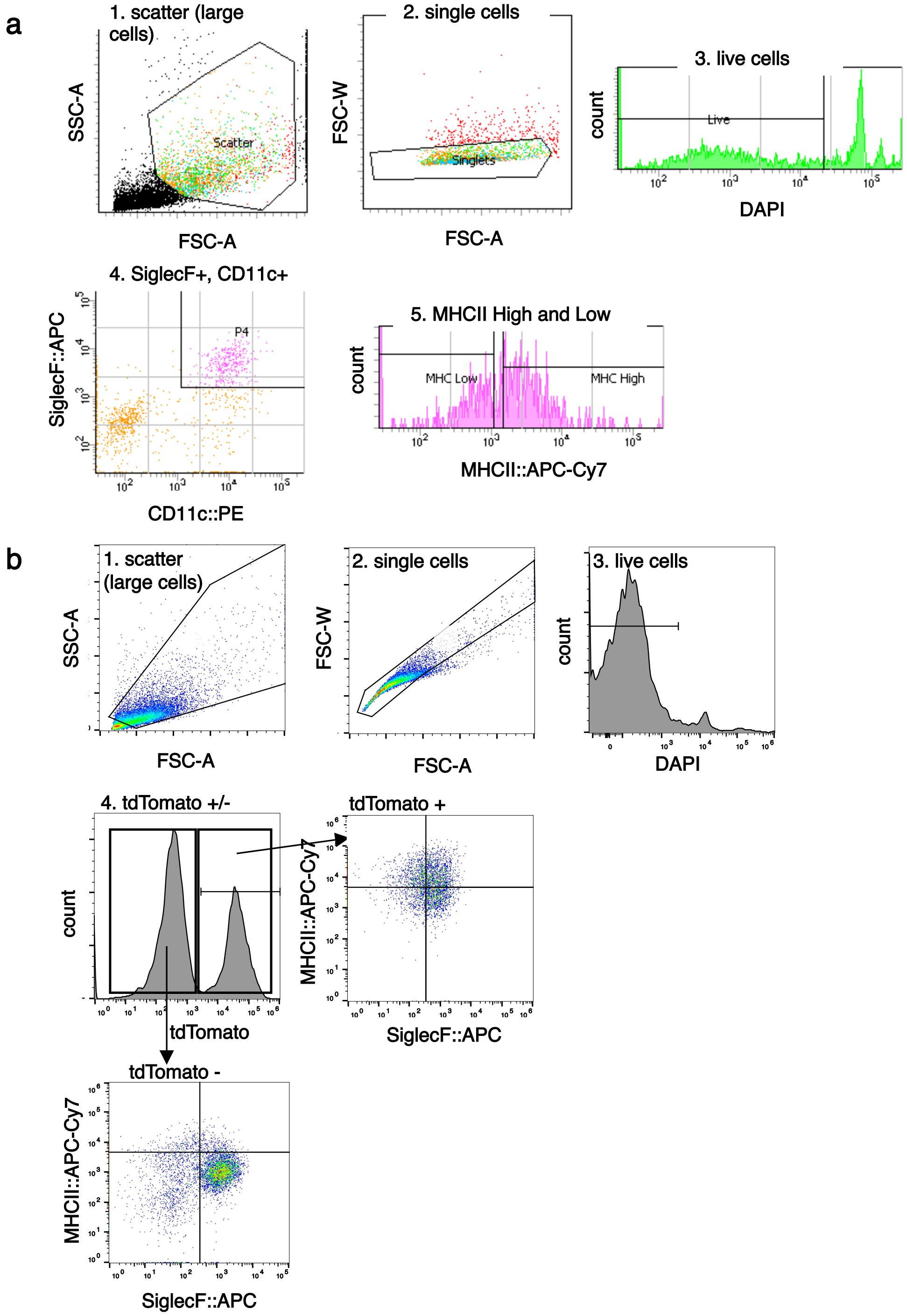

**Figure S6. Flow cytometric gating. a.** Cells were dissociated from lungs of WT mice 14 days after bleomycin injury and stained with SiglecF, CD11c, and MHCII antibodies. SiglecF+CD11c+ cells were sorted into MHCII-high and MHC-low populations, with the threshold defined by MHCII staining in an uninjured mouse. Representative data are shown. **b.** Lung cells were dissociated from CX3CR1CreERT2 / ROsa26$_{loxP}$STOP$_{loxP}$-TdTomato mice induced with tamoxifen 1 day prior to and during bleomycin injury and stained with SiglecF and MHCII. Representative data are shown.

#### Supplementary Figure 7

**a**

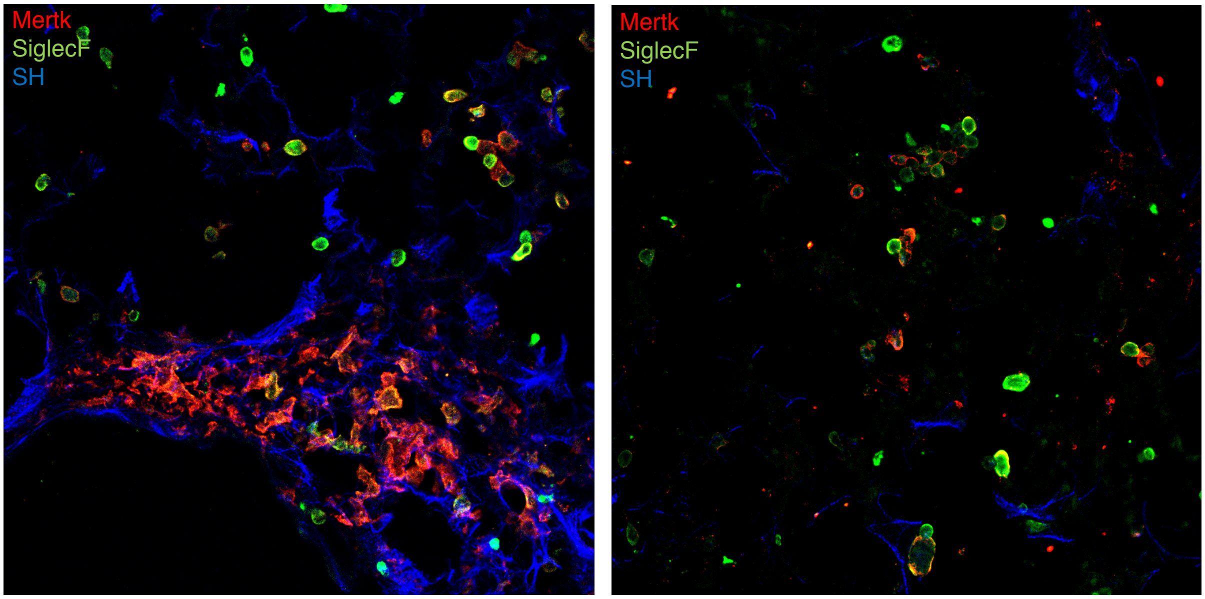

**b**

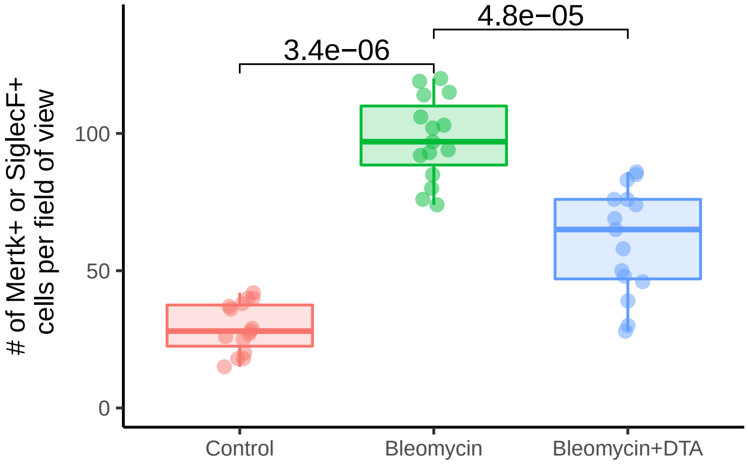

**Figure S7. Depletion of macrophage subpopulation during bleomycin fibrosis. a.** Fluorescence microscopy demonstrating diminished fibrotic scar, detected by 2-photon imaging of second harmonic (SH), and ablation of macrophages localized to scar by Mertk immunolabeling 21 days after bleomycin injury in tamoxifen-induced Cx3cr1CreERT2 x Rosa26$_{loxP}$STOP$_{loxP}$-Diptheria Toxin A mouse (right) compared with wild type control (left). **b.** Quantitation of total number of Mertk or SiglecF-expressing cells in 5 high power fields (n=3 mice per group). Wilcoxon rank sum test p-value is presented.

#### Supplementary Table 1

Table S1. Drop-seq batches.

| Set name, | Seq. Platform | Read length | Num. reads | Num. alinged reads | Num. cells | Num. >500 genes |
|---|---|---|---|---|---|---|
| Control_1 (X2) | Miseq | 25X75 | 56,367,089 | 70.97% | 2,000 | 706 |
| Control_2 | Hiseq2500 | 20X50 | 158,334,751 | 76.78% | 4,000 | 856 |
| Control_3 | Hiseq2500 | 20X50 | 112,676,785 | 81.80% | 6,000 | 662 |
| Control_4 | Hiseq4000 | 20X50 | 131,761,368 | 47.57% | 4,000 | 606 |
| Control_5 | Hiseq4000 | 20X50 | 119,391,037 | 60.66% | 4,000 | 492 |
| Control_6 | Hiseq2500 | 20X50 | 148,515,763 | 73.81% | 6,000 | 278 |
| Bleomycin_1 | Hiseq2500 | 20X50 | 115,586,230 | 77.38% | 4,000 | 1,552 |
| Bleomycin_2 | Hiseq2500 | 20X50 | 131,539,209 | 66.72% | 6,000 | 1,010 |
| Bleomycin_3 | Hiseq4000 | 20X50 | 235,055,677 | 68.81% | 6,000 | 2,204 |
| All controls | | | 727,046,793 | 68.56% | 26,000 | 3,600 |
| All bleomycin | | | 482,181,116 | 70.29% | 16,000 | 4,766 |

#### Supplementary Table 2

Table S2. Differentially expressed genes between C1 and C3.

| | P-value | Log 2FC | % of C1 | % of C3 | Adj. P-value |
|---|---|---|---|---|---|
| Chil3 | <1e-300 | 3.466798 | 0.994 | 0.304 | <1e-300 |
| 1100001g20rik | <1e-300 | 2.735837 | 0.848 | 0.091 | <1e-300 |
| Atp6v0d2 | <1e-300 | 2.372622 | 0.939 | 0.125 | <1e-300 |
| Cd9 | <1e-300 | 1.348099 | 0.998 | 0.801 | <1e-300 |
| S100a6 | <1e-300 | -1.792623 | 0.169 | 0.814 | <1e-300 |
| Plbd1 | <1e-300 | -1.894910 | 0.007 | 0.497 | <1e-300 |
| Mir5107 | <1e-300 | -1.937692 | 0.094 | 0.721 | <1e-300 |
| S100a4 | <1e-300 | -2.241013 | 0.133 | 0.852 | <1e-300 |
| Ccr2 | <1e-300 | -2.534832 | 0.095 | 0.784 | <1e-300 |
| Mgl2 | <1e-300 | -2.665418 | 0.008 | 0.475 | <1e-300 |
| C1qc | <1e-300 | -2.803449 | 0.019 | 0.693 | <1e-300 |
| H2-ab1 | <1e-300 | -2.901784 | 0.281 | 0.977 | <1e-300 |
| C1qa | <1e-300 | -2.916089 | 0.020 | 0.703 | <1e-300 |
| H2-aa | <1e-300 | -2.944505 | 0.211 | 0.970 | <1e-300 |
| C1qb | <1e-300 | -3.068720 | 0.034 | 0.816 | <1e-300 |
| H2-eb1 | <1e-300 | -3.243792 | 0.206 | 0.965 | <1e-300 |
| Chil4 | 7.90888491040646e-307 | 2.537923 | 0.808 | 0.028 | 1.09443149390205e-302 |
| Aif1 | 3.19901619542006e-289 | -1.694715 | 0.008 | 0.430 | 4.42679861122228e-285 |
| Srgn | 3.81536395648136e-270 | -1.502502 | 0.582 | 0.931 | 5.27970064297891e-266 |
| Il18 | 2.57986644164827e-255 | 1.767475 | 0.842 | 0.209 | 3.57001918195288e-251 |
| Ctsd | 1.13215934242929e-251 | 1.338009 | 0.956 | 0.531 | 1.56668209805366e-247 |
| Socs3 | 1.24309916394746e-245 | -1.612644 | 0.075 | 0.545 | 1.72020062307049e-241 |
| Cx3cr1 | 1.02341013376724e-241 | -1.567244 | 0.017 | 0.402 | 1.41619494310711e-237 |
| Cxcl16 | 4.12880896458557e-241 | -1.501350 | 0.060 | 0.512 | 5.71344584519351e-237 |
| Slpi | 1.54864297931837e-237 | 2.085565 | 0.810 | 0.171 | 2.14301215478076e-233 |
| Mmp12 | 4.09443364979244e-237 | -1.994885 | 0.056 | 0.501 | 5.66587728458278e-233 |
| Mcemp1 | 1.00419911579153e-232 | 1.511032 | 0.844 | 0.254 | 1.38961073643231e-228 |
| Cdc42ep3 | 1.51750523114026e-232 | 1.839832 | 0.763 | 0.120 | 2.0999237388519e-228 |
| Cd14 | 8.58679545916112e-231 | -1.264637 | 0.680 | 0.938 | 1.18824075563872e-226 |
| Ccl6 | 4.8613155307177e-229 | 1.278130 | 0.985 | 0.785 | 6.72708843140715e-225 |
| Ly86 | 8.12102386611689e-222 | -1.458767 | 0.087 | 0.540 | 1.12378728259326e-217 |
| Ear1 | 3.47158905304245e-221 | 1.830027 | 0.852 | 0.297 | 4.80398493160014e-217 |
| Mgst1 | 7.34258179925425e-217 | 1.363636 | 0.864 | 0.357 | 1.0160664693808e-212 |
| Ccl9 | 1.11663470499323e-216 | -1.566697 | 0.151 | 0.630 | 1.54519910476963e-212 |
| Cst3 | 2.68011592476351e-215 | -1.102656 | 0.954 | 0.985 | 3.70874441668775e-211 |
| Txn1 | 1.19048509496134e-211 | 1.067131 | 0.944 | 0.608 | 1.64739327440751e-207 |
| Cd52 | 6.42027574773743e-210 | -1.282794 | 0.512 | 0.876 | 8.88437757971905e-206 |
| Plet1 | 3.96151294627678e-209 | 1.729488 | 0.693 | 0.058 | 5.4819416150578e-205 |
| Sdc4 | 6.32185289490639e-204 | -1.492497 | 0.033 | 0.400 | 8.74818003597146e-200 |
| Junb | 1.19871101140209e-196 | -1.035783 | 0.700 | 0.945 | 1.65877629757821e-192 |
| Ctsk | 2.72549742918959e-187 | 1.878161 | 0.673 | 0.094 | 3.77154334251256e-183 |
| Fosb | 1.78344133939182e-185 | -1.296025 | 0.499 | 0.841 | 2.4679261254504e-181 |
| Il1b | 1.16521380971596e-176 | -1.677662 | 0.487 | 0.832 | 1.61242286988495e-172 |
| Car4 | 3.93277465115084e-170 | 2.203684 | 0.588 | 0.042 | 5.44217356226253e-166 |
| Hexb | 6.01841318716772e-168 | -1.265074 | 0.425 | 0.790 | 8.32828016840269e-164 |
| Abcg1 | 2.62827415093067e-165 | 1.324646 | 0.762 | 0.260 | 3.63700577005787e-161 |
| H2-dmb1 | 5.50609569227619e-164 | -1.241115 | 0.219 | 0.640 | 7.61933521897179e-160 |
| Mafb | 1.0347887685647e-162 | -1.505033 | 0.052 | 0.387 | 1.43194069793983e-158 |

| | P-value | Log 2FC | % of C1 | % of C3 | Adj. P-value |
|---|---|---|---|---|---|
| Plin2 | 1.10768485245912e-159 | 1.106772 | 0.838 | 0.392 | 1.53281429883293e-155 |
| Btg2 | 2.74718608057858e-159 | -1.200297 | 0.471 | 0.813 | 3.80155609830463e-155 |
| Ear10 | 7.26249012526558e-154 | 1.857297 | 0.703 | 0.197 | 1.00498338353425e-149 |
| Egr1 | 1.77993858634046e-152 | -1.432402 | 0.241 | 0.650 | 2.46307901577793e-148 |
| Marcks | 1.80940234676992e-152 | -1.306145 | 0.126 | 0.504 | 2.50385096746021e-148 |
| Plxdc2 | 3.26686663012579e-143 | -1.225561 | 0.087 | 0.429 | 4.52069004276806e-139 |
| Arl4c | 2.34860201246099e-138 | -1.182227 | 0.085 | 0.418 | 3.24999546484352e-134 |
| Krt79 | 1.21529743655194e-137 | 1.728516 | 0.486 | 0.015 | 1.68172859270057e-133 |
| Zfp36l1 | 6.65161853307967e-135 | -1.447980 | 0.119 | 0.470 | 9.20450972607565e-131 |
| Ltc4s | 1.66513563582204e-133 | 1.303861 | 0.542 | 0.060 | 2.30421469285054e-129 |
| Nfkbia | 9.32548888833289e-133 | -1.155190 | 0.440 | 0.770 | 1.29046115236751e-128 |
| Fpr2 | 5.02469306560165e-122 | 1.223014 | 0.542 | 0.077 | 6.95317026417956e-118 |
| Marco | 8.02808241540788e-121 | 2.226533 | 0.461 | 0.032 | 1.11092604464414e-116 |
| Pnpla8 | 2.06436965990703e-119 | 1.087071 | 0.689 | 0.255 | 2.85667473537934e-115 |
| Serpinb1a | 5.39836606612562e-114 | 1.381527 | 0.469 | 0.044 | 7.47025896230464e-110 |
| F7 | 1.95316410290726e-113 | 1.127582 | 0.642 | 0.195 | 2.70278848560306e-109 |
| Ccrl2 | 8.41872044137331e-111 | -1.195926 | 0.259 | 0.590 | 1.16498253467724e-106 |
| Kctd12 | 1.10794961536521e-108 | -1.088315 | 0.122 | 0.435 | 1.53318067774238e-104 |
| Adipor2 | 7.47013908782941e-108 | 1.072546 | 0.629 | 0.212 | 1.03371784697383e-103 |
| Fabp1 | 1.58663251510657e-106 | 2.210323 | 0.386 | 0.000 | 2.19558207440447e-102 |
| Gadd45b | 5.3185557834464e-106 | -1.057016 | 0.255 | 0.590 | 7.35981749313313e-102 |
| Fpr1 | 8.81726549273493e-105 | 1.262564 | 0.443 | 0.043 | 1.22013319888466e-100 |
| Krt19 | 1.17058119200157e-104 | 1.417590 | 0.392 | 0.008 | 1.61985025349177e-100 |
| Ppp1r15a | 2.93737811916475e-104 | -1.070439 | 0.097 | 0.388 | 4.06474384130018e-100 |
| Lilra5 | 1.07913273186667e-100 | 1.101697 | 0.525 | 0.117 | 1.4933038743571e-96 |
| Cfp | 1.29781646293635e-98 | -1.062722 | 0.250 | 0.562 | 1.79591842141132e-94 |
| Fabp4 | 2.27520559103527e-95 | 1.655911 | 0.369 | 0.010 | 3.1484294968746e-91 |
| Cd2 | 1.82291633248731e-92 | 1.078541 | 0.536 | 0.150 | 2.52255162089593e-88 |
| Pla2g7 | 1.16699283999901e-90 | -1.199363 | 0.195 | 0.488 | 1.61488469199063e-86 |
| Napsa | 2.91149644742703e-85 | -1.020722 | 0.213 | 0.502 | 4.02892878394953e-81 |
| Prkar2b | 1.12910050595557e-84 | 1.112273 | 0.428 | 0.077 | 1.56244928014132e-80 |
| Ier3 | 1.31318154110729e-80 | -1.189059 | 0.195 | 0.479 | 1.81718061658427e-76 |
| Gpnmb | 2.28081067880061e-74 | 1.213541 | 0.368 | 0.049 | 3.15618581732429e-70 |
| Cfh | 1.42287636261063e-66 | -1.167381 | 0.141 | 0.376 | 1.96897631058058e-62 |

#### Supplementary Table 3

Table S3. Mouse and Human orthologs of C1 genes

| Mouse.C1.genes | Human.C1.genes |
|---|---|
| Ctsk | CTSK |
| Chil4 | CHIA |
| Chil3 | CHIA |
| Cdc42ep3 | CDC42EP3 |
| Atp6v0d2 | ATP6V0D2 |
| Fabp4 | FABP4 |
| Ear10 | RNASE3 |
| Ear10 | RNASE2 |
| Ear1 | RNASE3 |
| Ear1 | RNASE2 |
| Marco | MARCO |
| Car4 | CA4 |
| Plet1 | PLET1 |
| Il18 | IL18 |
| Fabp1 | FABP1 |
| Krt19 | KRT19 |

#### Extended Data References

Altboum, Zeev, Yael Steuerman, Eyal David, Zohar Barnett-Itzhaki, Liran Valadarsky, Hadas Keren-Shaul, Tal Meningher, et al. 2014. "Digital cell quantification identifies global immune cell dynamics during influenza infection." *Molecular Systems Biology* 10 (January): 720. doi:10.1002/msb.134947.

Gibbings, Sophie L., Stacey M. Thomas, Shaikh M. Atif, Alexandra L. McCubbrey, A. Nicole Desch, Thomas Danhorn, Sonia M. Leach, et al. 2017. "Three Unique Interstitial Macrophages in the Murine Lung at Steady State." *American Journal of Respiratory Cell and Molecular Biology* 57 (1): 66–76. doi:10.1165/rcmb.2016-0361OC.

Gong, Ting, and Joseph D. Szustakowski. 2013. "DeconRNASeq: a statistical framework for deconvolution of heterogeneous tissue samples based on mRNA-Seq data." *Bioinformatics* 29 (8): 1083–5. doi:10.1093/bioinformatics/btt090.
