## Supplementary Materials for "Reference-based annotation of single cell transcriptomes identifies a profibrotic macrophage niche after tissue injury"

### SingleR 1.0 - Method overview and examples

#### Introduction

Recent advances in single cell RNA-seq (scRNA-seq) have enabled an unprecedented level of granularity in characterizing gene expression changes in disease models. Multiple single cell analysis methodologies have been developed to detect gene expression changes and to cluster cells by similarity of gene expression. However, the classification of clusters by cell type relies heavily on known marker genes, and the annotation of clusters is performed manually. This strategy suffers from subjectivity and limits adequate differentiation of closely related cell subsets. Here, we present *SingleR*, a novel computational method for unbiased cell type recognition of scRNA-seq. *SingleR* leverages reference transcriptomic datasets of pure cell types to infer the cell of origin of each of the single cells independently. *SingleR*'s annotations combined with Seurat, a processing and analysis package designed for scRNA-seq, provide a powerful tool for the investigation of scRNA-seq data. We developed an R package to generate annotated scRNA-seq objects that can then use the *SingleR* web tool for visualization and further analysis of the data – http://comphealth.ucsf.edu/SingleR.

Here we explain in details the *SingleR* pipeline and present examples of applying SingleR on publicly available mouse and human scRNA-seq datasets.

#### ***SingleR* specifications**

**Reference set:** A comprehensive transcriptomic dataset (microarray or RNA-seq) of pure cell types, preferably with multiple samples per cell type.

- **Mouse:** We processed and annotated two reference mouse datasets:
  - Immunological Genome Project (ImmGen): a collection of 830 microarray samples, which we classified to 20 main cell types and further annotated to 253 subtypes.
  - A dataset of 358 mouse RNA-seq samples annotated to 28 cell types. This dataset was collected, processed and shared, courtesy of Bérénice Benayoun. This data set is especially useful for brain-related samples.
- **Human:** For human datasets we use the following reference datasets:
  - Human Primary Cell Atlas (HPCA): a collection of Gene Expression Omnibus (GEO datasets), which contains 713 microarray samples classified to 38 main cell types and further annotated to 169 subtypes.

   - Blueprint+Encode: Blueprint Epigenomics, 144 RNA-seq pure immune samples annotated to 28 cell types. Encode: 115 RNA-seq pure stroma and immune samples annotated 17 cell types. Altogether, 259 samples with to 43 cell types.

- For specific applications, smaller datasets can be applicable (see the main paper for the macrophages example). *SingleR* is flexible to be used with any reference dataset.
- *SingleR* runs in two modes - considering all cell types and states, or aggregating cell states to main cell types. It is suggested to start the analysis by using 'main types' mode before diving deeper to all cell states.

**Single-cell set:** Single-cell RNA-seq dataset. It is a good practice to filter-out cells with non-sufficient genes identified and genes with non-sufficient expression across cells. In all examples below we filtered-out cells containing less than 500 genes, and considered genes which were found in at least one sample. However, we did not observe a significant decrease in performance when considering less stringent thresholds for the number of genes per cell.

**Annotation:** *SingleR* runs in two modes: (1) Single-cell: the annotation is performed for each single-cell independently. (2) Cluster: the annotation is performed on predefined clusters, where the expression of a cluster is the sum expression of all cells in the cluster. In addition, *SingleR* annotates cells/clusters to all reference cell types, or combines reference cell types to major cell types.

- **Step 1:** Spearman coefficient is calculated for single-cell expression with each of the samples in the reference dataset. The correlation analysis is performed only on variable genes in the reference dataset. In all examples below we chose variable genes using a cutoff on the standard deviation; however other methods can be applied.

```
load (file.path(path,'GSE74923.RData'))
load (file.path(path,'references/Immgen.RData'))
SingleR.DrawScatter(sc_data = singler$seurat@data,cell_id = 10,
                    ref = immgen, sample_id = 232)
```

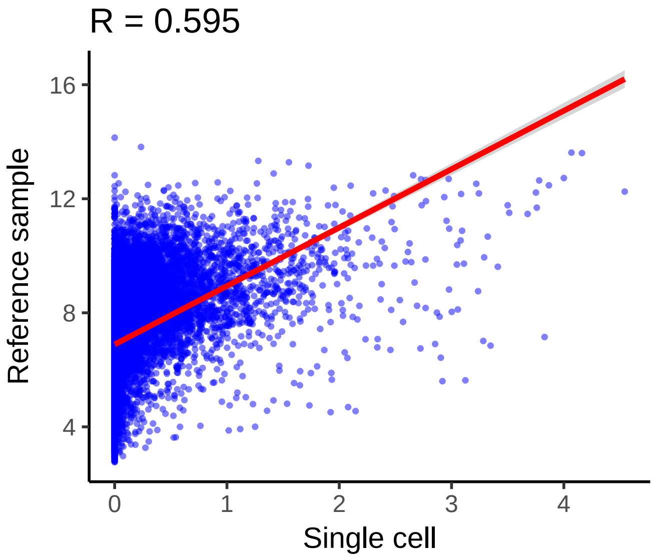

- **Variable genes:** *SingleR* supports to modes for choosing the variable genes in the reference dataset.

1. 'sd' - genes with a standard deviation across all samples in the reference dataset over a threshold. We choose thresholds such that we start with 3000-4000 genes.
2. 'de' - top N genes that have a higher median expression in a cell type compared to each other cell type. We use a varying N, depending on the number of cell types used in the analysis. More details can be found in the *SingleR* code. This mode was used in all examples below and in the web tool.

- **Step 2:** Multiple correlation coefficients per cell types are aggregated to provide a single value per cell type per single-cell. In the examples below we use the 80% percentile of correlation values.

```
# for visualization purposes we only present a subset of cell types (defined in labels.use)
out = SingleR.DrawBoxPlot(sc_data = singler$seurat@data,cell_id = 10,
                          ref = immgen,main_types = T,
labels.use=c('B cells','T cells','DC','Macrophages','Monocytes','NK cells',
             'Mast cells','Neutrophils','Fibroblasts','Endothelial cells'))
```

```
## [1] "Number of genes with SD>0.9: 2307"
## [1] "Number of cells: 2"
```

```
print(out$plot)
```

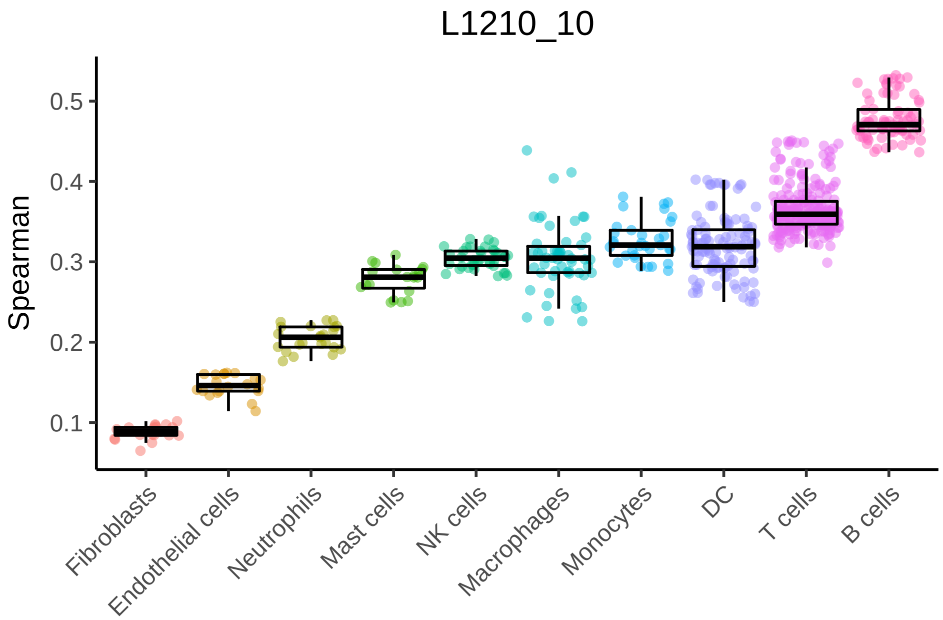

- **Step 3:** In this step *SingleR* reruns the correlation analysis, but only for the top cell types from step 2. The analysis is performed only on **variable genes between these cell types**. The lowest value cell type is removed (or a margin of more than 0.05 from top value), and then this step is repeated until only two cell types remain. The cell type corresponding to the top value after the last run is assigned to the single-cell.

The open source code is available in the *SingleR* Github repository https://github.com/dviraran/SingleR.

#### Running *SingleR*

We provide wrapper functions to create *SingleR* objects. **CreateSinglerSeuratObject** generates a SingleR object holding a Seurat object. **CreateSinglerObject** generates solely a *SingleR* object and metadata can be added separately. Both functions run *SingleR* with the relevant reference datasets described above. Click on 'code' to see an example.

```
# Generating SingleR+Seurat object
singler = CreateSinglerSeuratObject(counts = 'GSE74923_series_matrix.txt',
       annot = 'GSE74923_types.txt', project.name = 'GSE74923', min.genes = 500,
       min.cells = 2, npca = 10, regress.out ='nUMI', technology= 'C1', species = 'Mouse',
       citation = 'Kimmerling et al. 2016',reduce.file.size = T, variable.genes = 'de'
       normalize.gene.length = T)
```

```
# Generating SingleR object without Seurat. To use this SingleR visualization functions
# it is necessary to add coordinates for each cell in the field 'singler$meta.data$xy'
singler = CreateSinglerObject(counts = 'GSE74923_series_matrix.txt',
       annot = 'GSE74923_types.txt', project.name = 'GSE74923', min.genes = 500,
       technology= 'C1', species = 'Mouse',
       citation = 'Kimmerling et al. 2016',reduce.file.size = T, variable.genes = 'de'
       normalize.gene.length = T)
```

After reading the files with the counts and the annotations, the function creates a Seurat object using the wrapper function *SingleR.CreateSeurat* (only if using the first function). Next, if the scRNA-seq data is full-length the counts are normalized to gene length (TPM). Then, the *SingleR.CreateObject* is called for each reference data set. Finally, *calculateSignatures* is called to create reference datasets.

The resulting object is a list with the following fields:

- seurat - the Seurat object (only if using the relevant function). Using the reduce.file.size switch will remove the raw data from the object.
- singler - another list, with data for the singler annotations with each reference dataset. Each element in the list is a list containing the following fields:
  - SingleR.single - a list for the results by single cells for all cell types.
  - SingleR.single.main - a list for the results by single cells only for main cell types.
  - SingleR.cluster - a list for the results by clusters for all cell types.
  - SingleR.cluster.main - a list for the results by clusters only for main cell types.
- Each of those elements contains the following fields (among others):
  - scores - a matrix of the aggregated scores. Row for each single cell (or cluster), column for each cell type.
  - labels - the fine-tuned annotated labels.
  - labels1 - the annotated labels, without fine-tuning (after one round).
- signatures - a data frame with ssGSEA scores for prespecified signatures.
- meta.data - a list containing the project name, original identities and the coordinates.

#### Benchmarking *SingleR*

In the examples below we use the Seurat package to process the scRNA-seq data and perform the t-SNE analysis. All visualizations are readily available through the *SingleR* web tool – http://comphealth.ucsf.edu/SingleR. The web app allows to view the data and interactively analyze it.

##### Example 1: GSE74923 – Kimmerling et al. Nature Communications (2016)

A data set that was created to test the C1 platform. 189 single-cell mouse cell lines were analyzed using C1: 86 L1210 cells, mouse lymphocytic leukemia cells, and 103 mouse CD8+ T-cells.

First, we look at the t-SNE plot colored by the original identities:

```
# singler$singler[[1]] is the annotations obtained by using ImmGen dataset as reference.
# singler$singler[[2]] is based on the Mouse-RNAseq datasets.
out = SingleR.PlotTsne(singler$singler[[1]]$SingleR.single,
      singler$meta.data$xy, do.label = FALSE, do.letters = F,
      labels=singler$meta.data$orig.ident,label.size = 4,
      dot.size = 3)
out$p
```

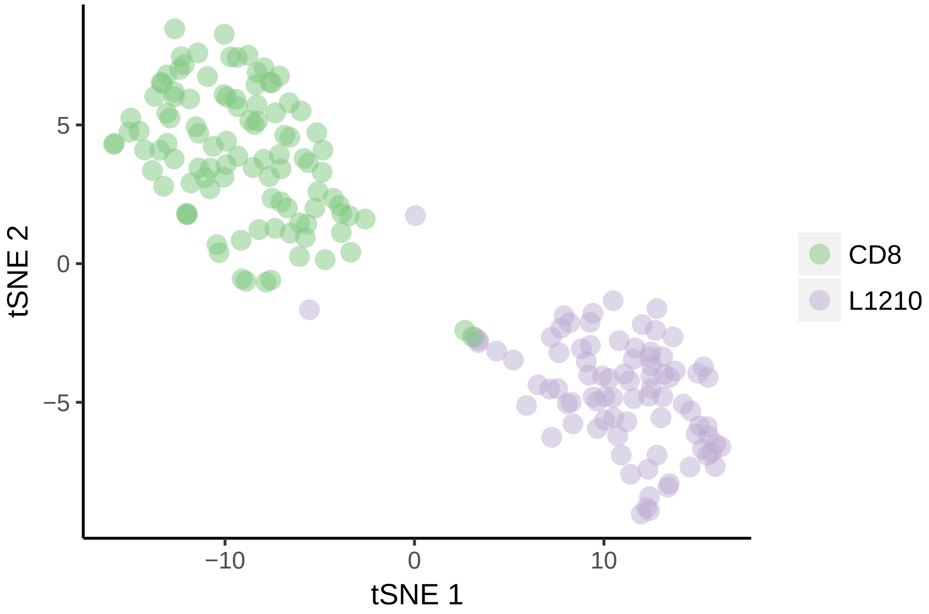

We can then observe the classification by a heatmap of the aggregated scores. These scores are before fine-tuning. We can view this heatmap by the main cell types:

```
SingleR.DrawHeatmap(singler$singler[[1]]$SingleR.single.main, top.n = Inf,
                    clusters = singler$meta.data$orig.ident)
```

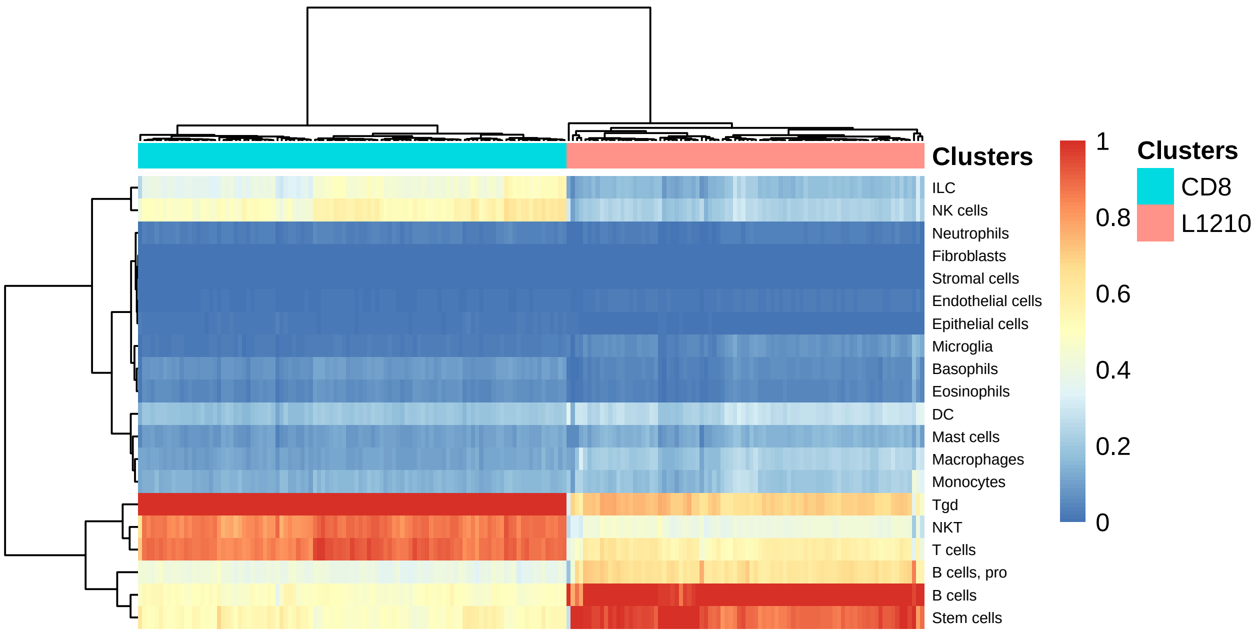

Or by all cell types (showing the top 50 cell types):

```
SingleR.DrawHeatmap(singler$singler[[1]]$SingleR.single, top.n = 50,
                    clusters = singler$meta.data$orig.ident)
```

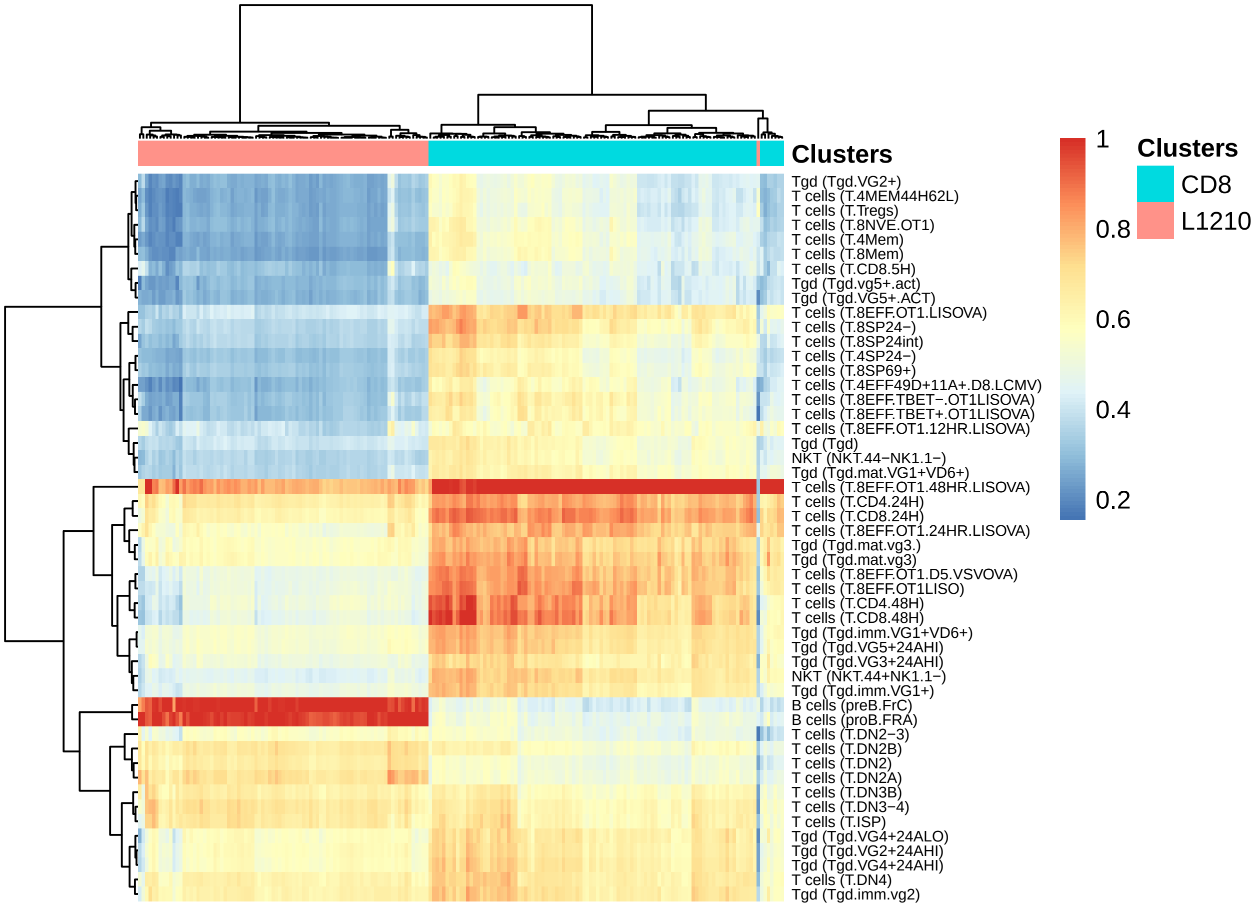

We can see that the L1210 cells were classified strongly to 3 types of B-cells progenitors. We can see that the CD8 cells were mostly correlated with a specific activation of effector CD8+ T-cells. Interestingly, there is one cell that seems to be correlated with both pro B-cells and CD8+ T-cells, suggesting that this a doublet.

Another interesting application of this heatmap is the ability to cluster the cells, not by their gene expression profile, but by their similarity to all cell types in the database. We use this clustering technique in the manuscript (Figure 2b).

Note: this view may be misleading, since each column is normalized to a score between 0 and 1 (which also depends on the cell types included in the view). Thus, a single cell may have low correlations with multiple cell types, and none of them are the accurate cell type, but in the heatmap they will all be red. Without normalization the data looks like this:

```
SingleR.DrawHeatmap(singler$singler[[1]]$SingleR.single,top.n = 50,
        normalize = F,clusters = singler$meta.data$orig.ident)
```

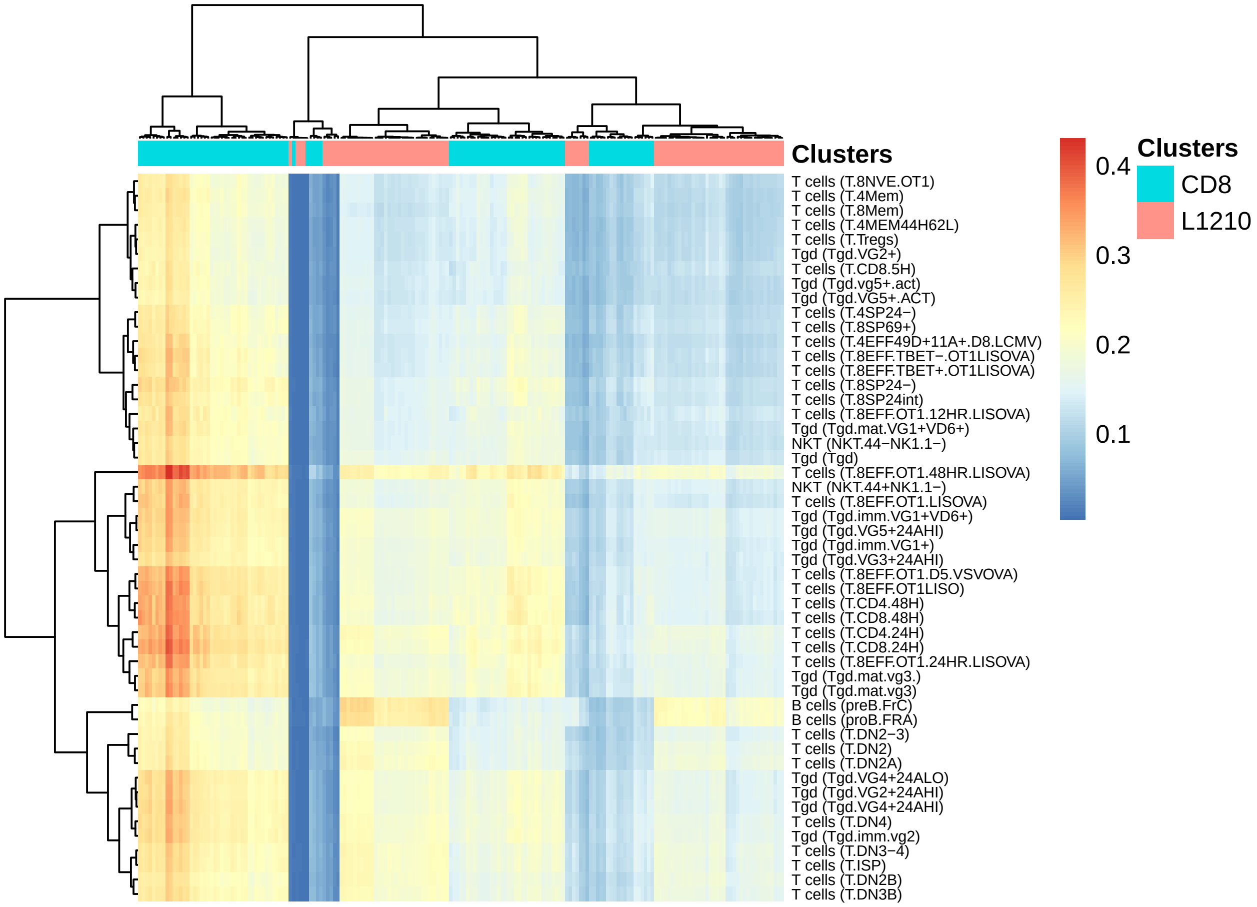

Next, we can use the fine-tuned labels to color the t-SNE plot:

```
out = SingleR.PlotTsne(singler$singler[[1]]$SingleR.single,
        singler$meta.data$xy,do.label=FALSE,
        do.letters = F,labels = singler$singler[[1]]$SingleR.single$labels,
        label.size = 4, dot.size = 3)
out$p
```

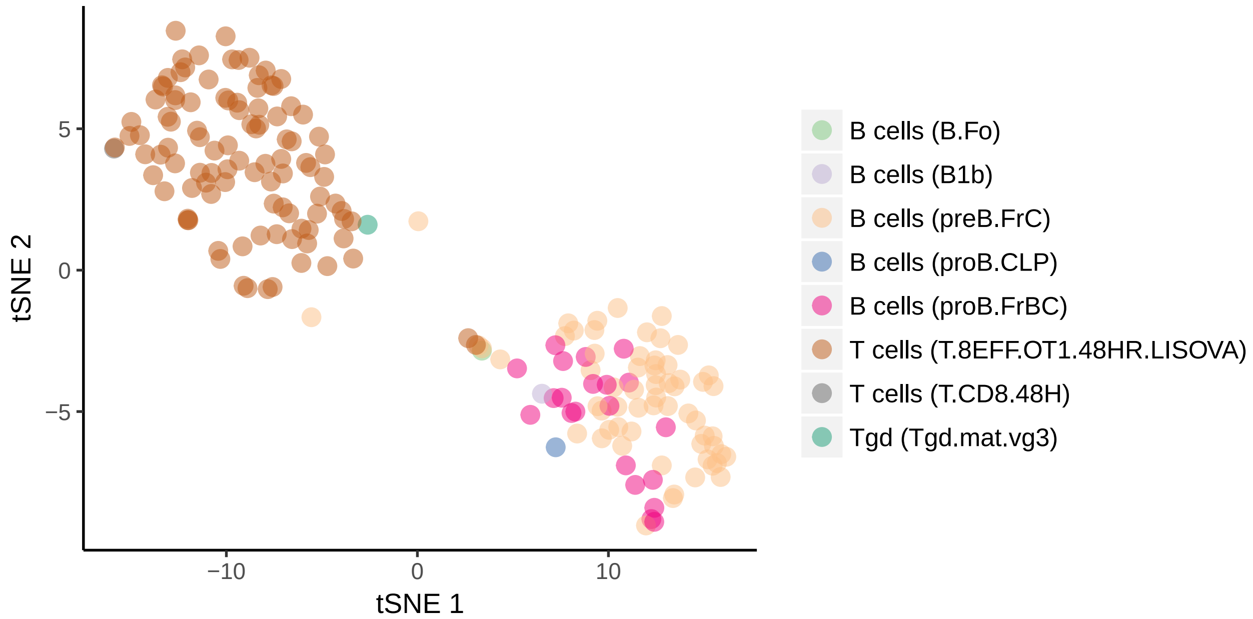

We can see that *SingleR* correctly annotated all the L1210 as types of B-cells, almost exclusively as B-cells progenitors. On the other hand, all CD8 cells were correctly annotated to CD8+ T-cells. It is important to remember that there are 253 types that *SingleR* could have chosen from, but it correctly chose the most relevant cell types. Interestingly, the tSNE plot incorrectly positioned cells in the wrong cluster, but *SingleR* was not affected by this.

Finally, we can also view the labeling as a table compared to the original identities:

```
kable(table(singler$singler[[1]]$SingleR.single$labels,singler$meta.data$orig.ident))
```

| | CD8 | L1210 |
|---|---|---|
| B cells (B.Fo) | 0 | 1 |
| B cells (B1b) | 0 | 1 |
| B cells (preB.FrC) | 0 | 62 |
| B cells (proB.CLP) | 0 | 1 |
| B cells (proB.FrBC) | 0 | 21 |
| T cells (T.8EFF.OT1.48HR.LISOVA) | 101 | 0 |
| T cells (T.CD8.48H) | 1 | 0 |
| Tgd (Tgd.mat.vg3) | 1 | 0 |

###### Example 2: GSE48968 – Shalek et al. Nature (2014)

In the main manuscript we described our analysis to a dataset produced by Hashimshony et al. (2016), which contains scRNA-seq of fibroblasts and bone-marrow derived dendritic cells (BMDCs). We showed that 33 of those 48 cells were actually macrophages, in accordance with a Helft et al. (2015). Here we reanalyzed a seminal study in scRNA-seq, which performed one of the earliest large-scale analyses of single-cells. Using the Smart-Seq protocol the authors analyzed 1775 single mouse BMDCs. The paper described in detail different dendritic cells (DCs) activation states that have been observed in the data, without any mention of macrophages.

Running Seurat with the same filters as described above produced the following t-SNE plot:

```
load (file.path(path,'GSE48968.RData'))
out = SingleR.PlotTsne(singler$singler[[1]]$SingleR.single,
```

```
        singler$meta.data$xy, do.label = FALSE,
        do.letters = F,labels = singler$meta.data$orig.ident,
        dot.size = 1.5)
out$p
```

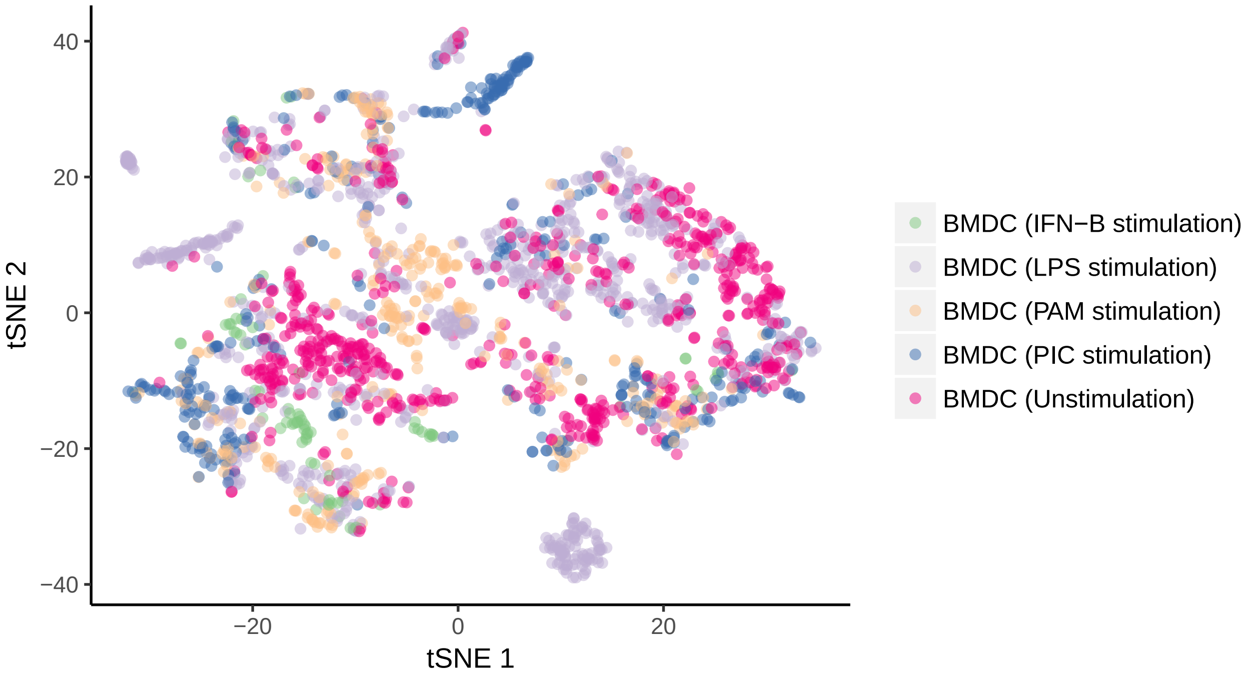

And the *SingleR* annotations (main cell types):

```
out = SingleR.PlotTsne(singler$singler[[1]]$SingleR.single.main,
        singler$meta.data$xy, do.label = FALSE,
        do.letters = F, dot.size = 2)
out$p
```

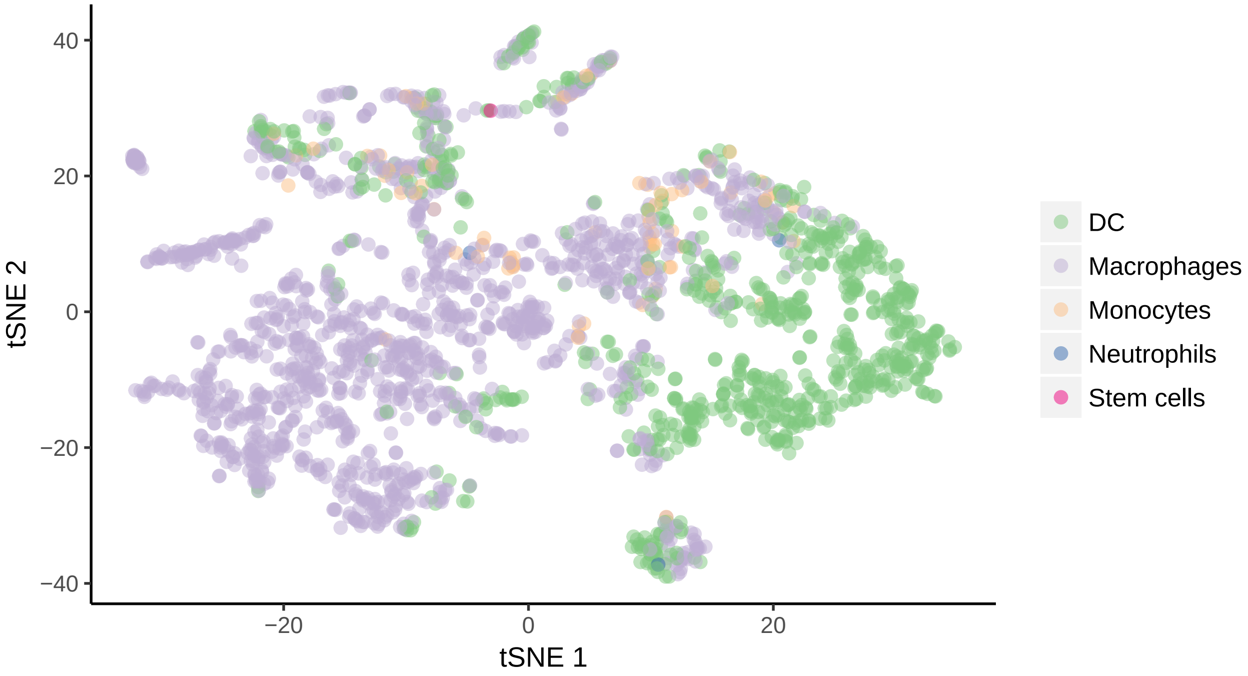

*SingleR* mapped all the left side to macrophages, top right to monocytes, and only the bottom right to dendritic cells.

In a table:

```
kable(table(singler$meta.data$orig.ident,singler$singler[[1]]$SingleR.single.main$labels))
```

| | DC | Macrophages | Monocytes | Neutrophils | Stem cells |
|---|---|---|---|---|---|
| BMDC (IFN-B stimulation) | 13 | 70 | 0 | 0 | 0 |
| BMDC (LPS stimulation) | 264 | 562 | 28 | 2 | 0 |
| BMDC (PAM stimulation) | 71 | 246 | 28 | 0 | 0 |
| BMDC (PIC stimulation) | 127 | 230 | 11 | 0 | 1 |
| BMDC (Unstimulation) | 343 | 303 | 11 | 1 | 0 |

As in the main manuscript, we employed the dataset produced by Helft et al. (2015) (downloaded from GEO accession: GSE62361), which analyzed by microarray the expression profiles of GM-DCs and GM-Macs. We select the top 50 deferentially expressed genes of each cell type, and present a heatmap of their expression in the single cells:

```
gse62631.de <- read.table(file.path(path,'GSE62361_DE.txt'),
                          header=TRUE, sep="\t", row.names=1, as.is=TRUE)
bmdc.genes = gse62631.de[intersect(rownames(gse62631.de),
                                   rownames(singler$seurat@data)),'Group',drop=F]
d = t(scale(t(as.matrix(singler$seurat@data[rownames(bmdc.genes),]))))
d[d>2]=2;d[d< -2]=-2
annotation_col = data.frame(Annotation=singler$singler[[1]]$SingleR.single.main$labels)
pheatmap(d[order(bmdc.genes$Group),
         order(singler$singler[[1]]$SingleR.single.main$labels)],
         cluster_cols = F,cluster_rows = F,clustering_method='ward.D',
         border_color = NA,
         annotation_col=annotation_col,
         annotation_row = bmdc.genes,show_colnames=F,show_rownames=F)
```

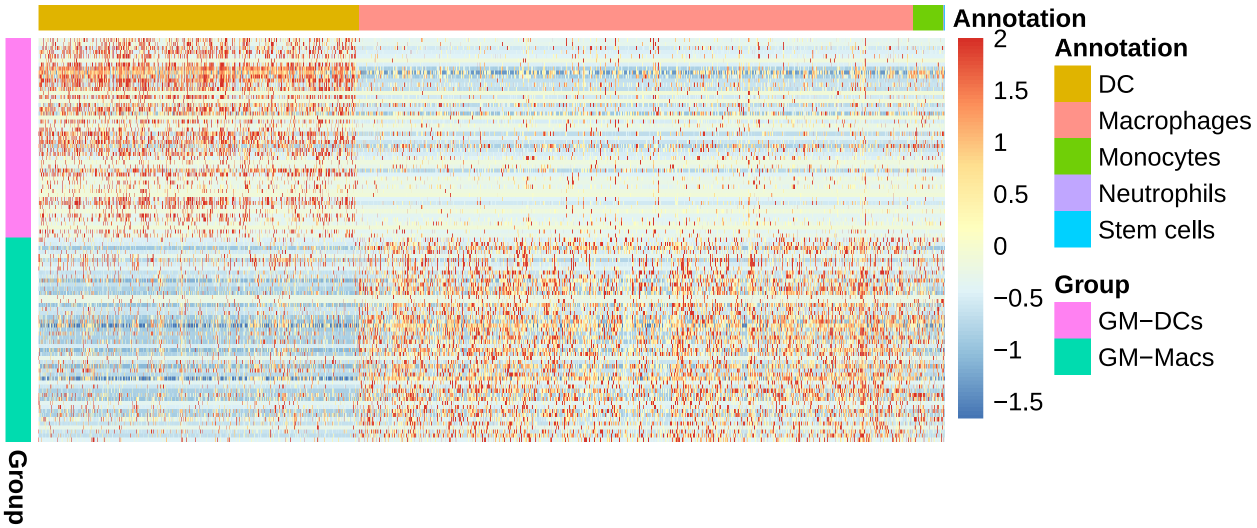

We can see that *SingleR* correctly identified the DCs and the macrophages populations. Interestingly, the cells annotated as monocytes seem to contain markers of both GM-Macs and GM-DCs, suggesting that those are earlier progenitors, which have not yet committed to the DC or Macrophage lineage.

In accordance with Helft et al. this analysis urges the reevaluation of studies to characterize DCs based on

BMDC. It also emphasizes how using *SingleR* can assist in resolving the cellular heterogeneity.

###### Example 3: 10X datasets – Zheng et al. Nature Communications (2017)

We also applied *SingleR* to human datasets. Using the 10X platform, Zheng et al. produced >100K single cells from sorted immune and cell lines populations. We obtained this data from https://support.10xgenomics.com/single-cell-gene-expression/datasets, and processed it with the *SingleR* pipeline. To reduce computation times and to make the analyses simpler, we randomly selected 200 cells with >500 non-zero genes from 10 immune populations.

```
load (file.path(path,'10x (Zheng) - 2000cells.RData'))
out = SingleR.PlotTsne(singler$singler[[1]]$SingleR.single,
            singler$meta.data$xy,do.label = FALSE,
            do.letters = F,labels = singler$meta.data$orig.ident,
            dot.size = 2)
out$p
```

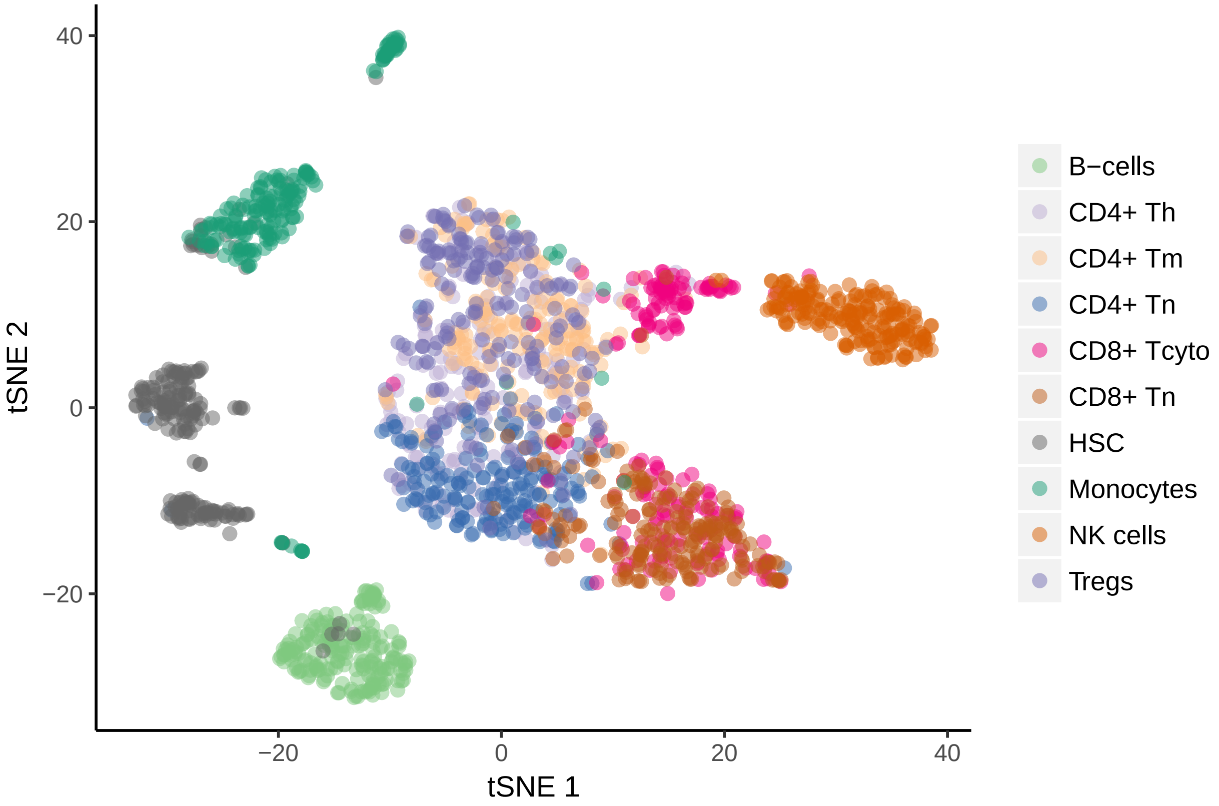

The tSNE plots allows to distinguish most cell types, but the T-cells subsets are all blurred together.

We first look at the Seurat clustering:

```
out = SingleR.PlotTsne(singler$singler[[1]]$SingleR.single,
            singler$meta.data$xy,do.label = T,
            do.letters = F,labels=singler$seurat@ident,
            dot.size = 2,label.size = 4)
out$p
```

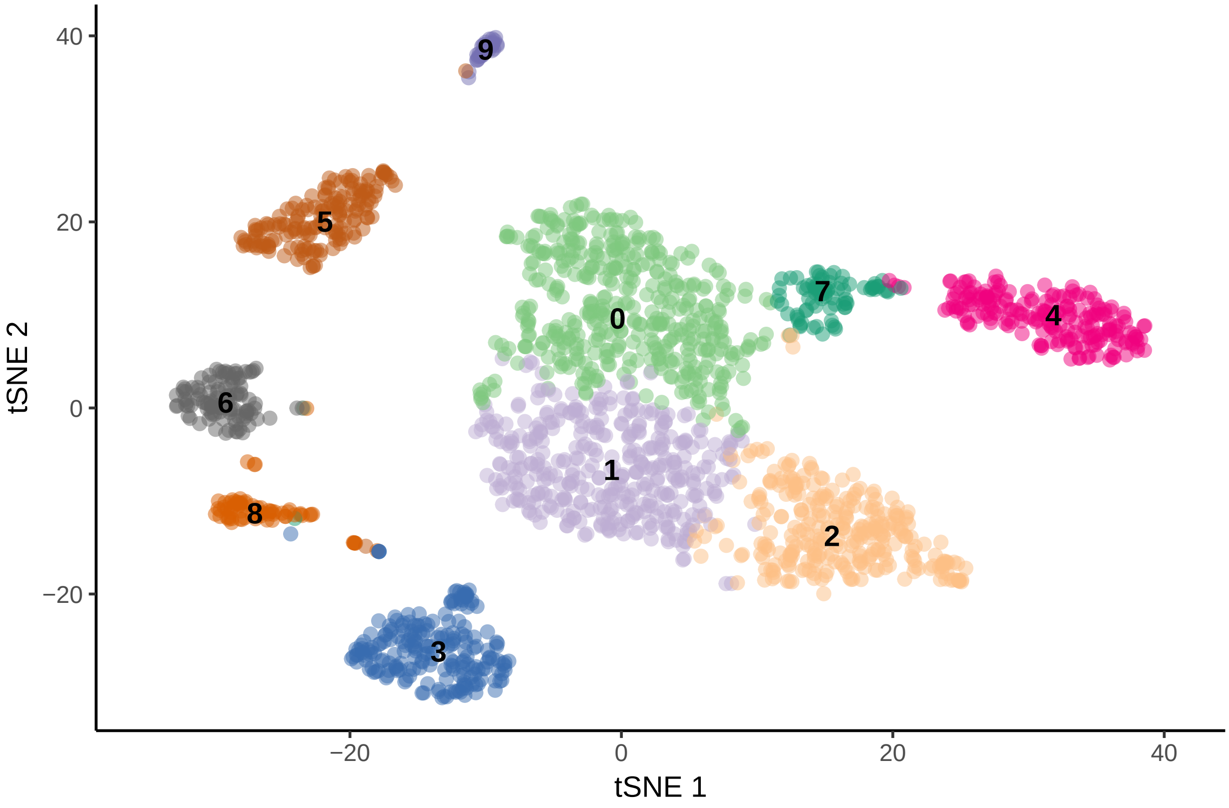

```
kable(table(singler$meta.data$orig.ident,singler$seurat@ident))
```

| | 0 | 1 | 2 | 3 | 4 | 5 | 6 | 7 | 8 | 9 |
|---|---|---|---|---|---|---|---|---|---|---|
| B-cells | 0 | 0 | 0 | 200 | 0 | 0 | 0 | 0 | 0 | 0 |
| CD4+ Th | 96 | 101 | 0 | 0 | 0 | 0 | 0 | 3 | 0 | 0 |
| CD4+ Tm | 167 | 24 | 6 | 0 | 0 | 0 | 0 | 3 | 0 | 0 |
| CD4+ Tn | 1 | 192 | 4 | 0 | 0 | 0 | 1 | 0 | 1 | 1 |
| CD8+ Tcyto | 7 | 9 | 104 | 0 | 6 | 0 | 0 | 74 | 0 | 0 |
| CD8+ Tn | 1 | 21 | 177 | 0 | 0 | 0 | 0 | 1 | 0 | 0 |
| HSC | 1 | 0 | 0 | 6 | 0 | 20 | 104 | 1 | 67 | 1 |
| Monocytes | 6 | 2 | 1 | 4 | 0 | 149 | 0 | 0 | 4 | 34 |
| NK cells | 0 | 0 | 0 | 0 | 199 | 0 | 0 | 1 | 0 | 0 |
| Tregs | 162 | 38 | 0 | 0 | 0 | 0 | 0 | 0 | 0 | 0 |

We can see that Seurat performs relatively well; however, regulatory T-cells are completely dissolved in the memory T-cells cluster.

*SingleR* using Blueprint+ENCODE (BE) as reference produced the following annotations before fine-tuning:

```
# Note the use of the second tiem in the the singler$singler list to use the Blueprint+ENCODE reference
# use singler$singler[[i]]$about for meta-data on the reference.
SingleR.DrawHeatmap(singler$singler[[2]]$SingleR.single,top.n=Inf,
                    clusters = singler$meta.data$orig.ident)
```

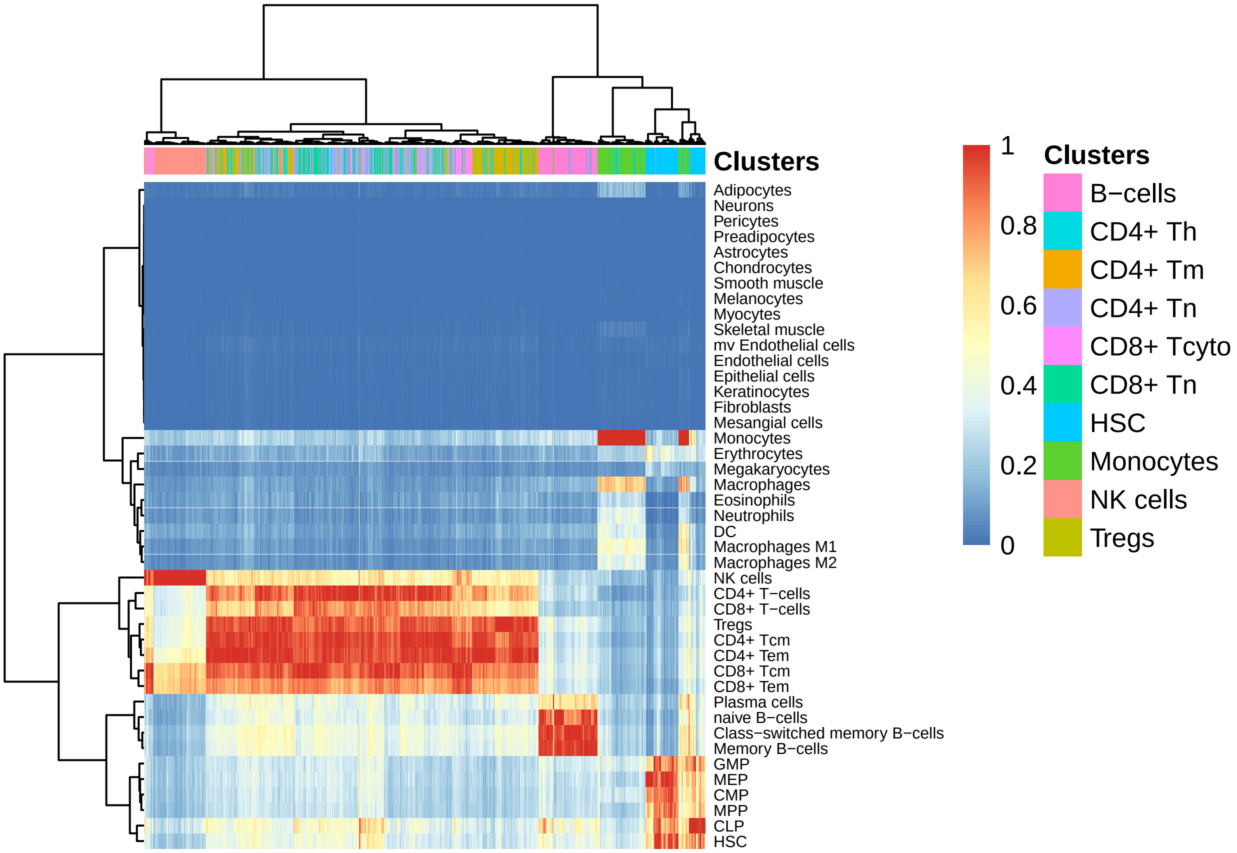

We can see that before fine-tuning, there is strong blurring between T-cells states, which cannot be distinguished.

However, with fine-tuning we obtain the following annotations:

```
out = SingleR.PlotTsne(singler$singler[[2]]$SingleR.single,
          singler$meta.data$xy,do.label=FALSE,
          do.letters =F,labels=singler$singler[[2]]$SingleR.single$labels,
          dot.size = 1.5, font.size = 6)
out$p
```

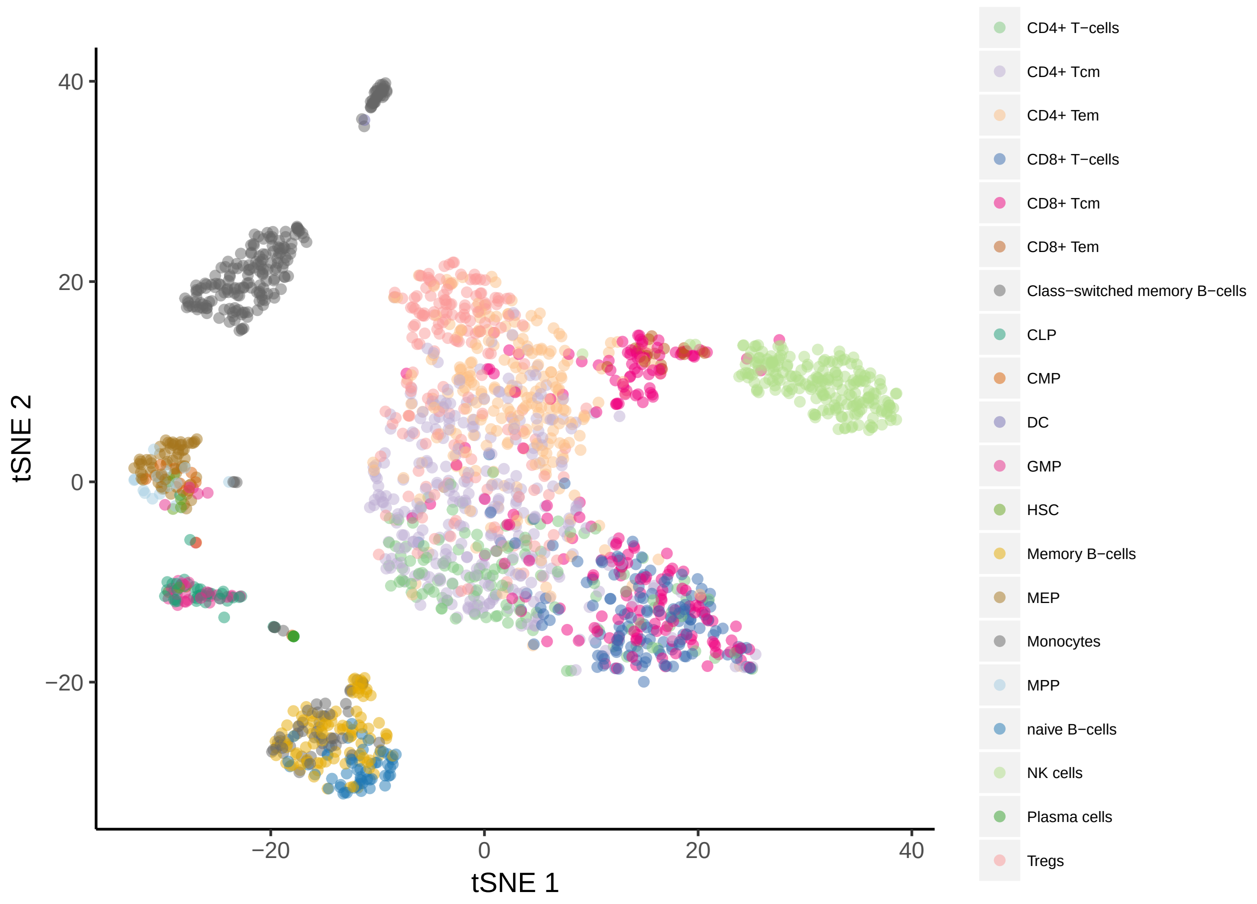

```
k = kable(table(singler$singler[[2]]$SingleR.single$labels,
          singler$meta.data$orig.ident), "latex")
kable_styling(k,font_size=6,bootstrap_options = "striped",
              full_width = F)
```

| | B-cells | CD4+ Th | CD4+ Tm | CD4+ Tn | CD8+ Tcyto | CD8+ Tn | HSC | Monocytes | NK cells | Tregs |
|---|---|---|---|---|---|---|---|---|---|---|
| CD4+ T-cells | 0 | 23 | 1 | 68 | 15 | 22 | 0 | 0 | 0 | 0 |
| CD4+ Tcm | 0 | 82 | 46 | 108 | 14 | 11 | 0 | 0 | 0 | 47 |
| CD4+ Tem | 0 | 57 | 97 | 7 | 12 | 3 | 0 | 6 | 0 | 59 |
| CD8+ T-cells | 0 | 1 | 0 | 1 | 38 | 91 | 0 | 2 | 0 | 0 |
| CD8+ Tcm | 0 | 9 | 9 | 4 | 105 | 72 | 0 | 0 | 0 | 4 |
| CD8+ Tem | 0 | 2 | 1 | 0 | 14 | 0 | 0 | 0 | 0 | 0 |
| Class-switched memory B-cells | 46 | 0 | 0 | 0 | 0 | 0 | 1 | 0 | 0 | 0 |
| CLP | 0 | 0 | 0 | 0 | 0 | 0 | 39 | 1 | 0 | 0 |
| CMP | 0 | 0 | 0 | 1 | 0 | 0 | 14 | 1 | 0 | 0 |
| DC | 0 | 0 | 0 | 0 | 0 | 0 | 0 | 1 | 0 | 0 |
| GMP | 0 | 0 | 0 | 0 | 0 | 0 | 29 | 0 | 0 | 0 |
| HSC | 0 | 0 | 0 | 0 | 0 | 0 | 7 | 0 | 0 | 0 |
| Memory B-cells | 97 | 0 | 0 | 0 | 0 | 0 | 4 | 0 | 0 | 0 |
| MEP | 0 | 0 | 0 | 0 | 0 | 0 | 62 | 0 | 0 | 0 |
| Monocytes | 0 | 0 | 0 | 1 | 0 | 0 | 25 | 184 | 0 | 0 |
| MPP | 0 | 0 | 0 | 1 | 0 | 0 | 19 | 0 | 0 | 0 |
| naive B-cells | 57 | 0 | 0 | 0 | 0 | 0 | 0 | 0 | 0 | 0 |
| NK cells | 0 | 0 | 0 | 0 | 0 | 0 | 0 | 1 | 200 | 0 |
| Plasma cells | 0 | 0 | 0 | 0 | 0 | 0 | 0 | 4 | 0 | 0 |
| Tregs | 0 | 26 | 46 | 9 | 2 | 1 | 0 | 0 | 0 | 90 |

By observing the colors we can see that the CD4+ T-cell cluster can roughly be divided in to 4 states, from naive CD4+ T-cells on the bottom (green), to central memory and effector memory CD4+ T-cells in the middle (purple and orange) and Tregs in the top (pink), in accordance with the original identities.

While it is not perfect, it provides us a much more granular view of the cell states without the need to go over many markers that might not be in the data at all, and thus whose interpretation is sometimes confusing:

```
df = data.frame(x=singler$meta.data$xy[,1],
                y=singler$meta.data$xy[,2],
                t(as.matrix(singler$seurat@data[c('CD3E','CD4','CD8A',
                        'CCR7','GZMA','GNLY','MS4A1','CD14','CD34'),])))
df = melt(df,id.vars = c('x','y'))
ggplot(df,aes(x=x,y=y,color=value)) +
  geom_point(size=0.3)+scale_color_gradient(low="gray", high="blue") +
  facet_wrap(~variable,ncol=3) +theme_classic()+xlab('')+ylab('')
```

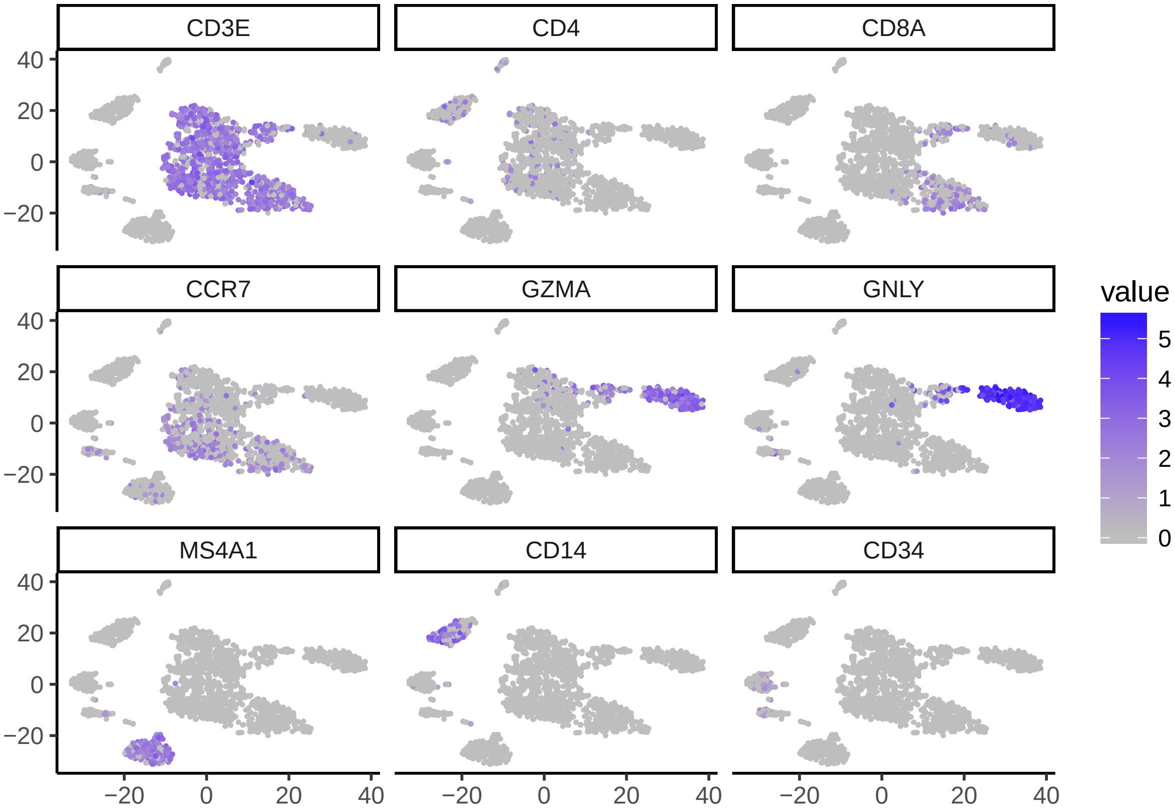

Interestingly, *SingleR* suggests a more granular view of the B-cell cluster, splitting it to naive and memory B-cells.

Finally, we compared our method to two other classification methods. Kang et al., Nature Biotechnology (2017) used sets of markers learned from scRNA-seq PBMCs (from Zheng et al.) to correlate each single cell:

```
out = SingleR.PlotTsne(singler$singler[[2]]$SingleR.single,
            singler$meta.data$xy,do.label=F,
            do.letters =F,labels=singler$other[,'Kang'],
            dot.size = 1.3)
out$p
```

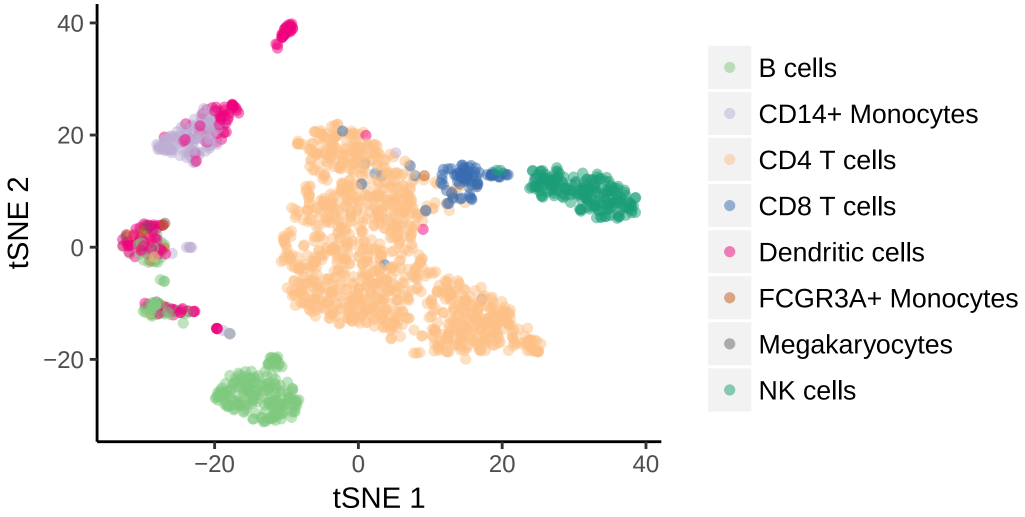

We can see that this methods has limited usability.

A bulk reference-based method by Li et. al, Nature Genetics, (2017) used a reference-based approach (but without fine-tuning).

```
out = SingleR.PlotTsne(singler$singler[[2]]$SingleR.single,
          singler$meta.data$xy,do.label=F,
          do.letters =F,labels=singler$other[,'RCA'],
          dot.size = 1.3,font.size=5)
out$p
```

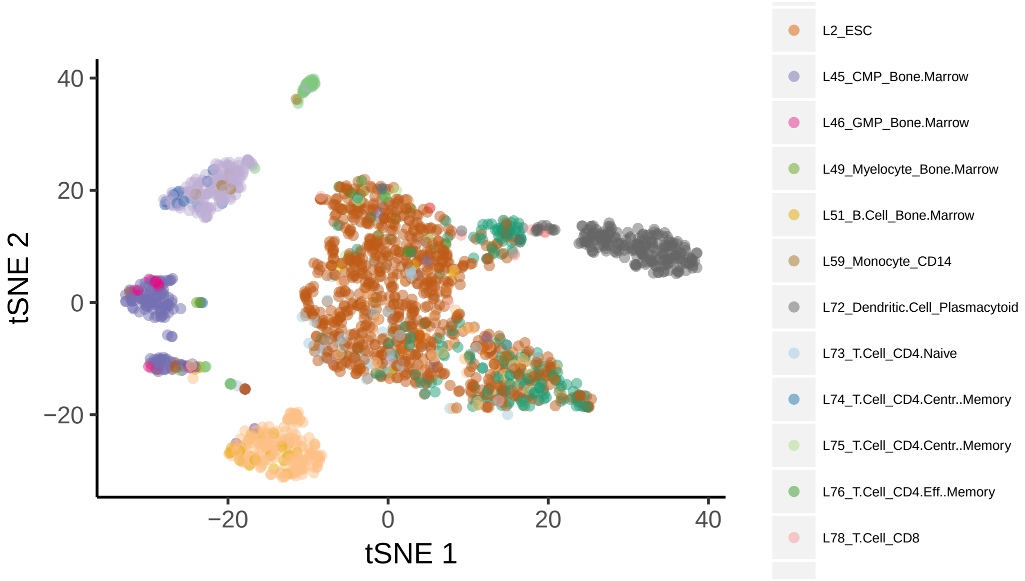

Here we can see that the microrray reference is not able to distinguish CD4+ and CD8+ T-cells, and without fine-tuning results it does not provide a sufficient solution for annotation.

*SingleR* also allows to detect rare events. For example, lets take a deeper look at the sorted Monocytes:

```
monocytes = SingleR.Subset(singler,singler$meta.data$orig.ident=='Monocytes')
out = SingleR.PlotTsne(monocytes$singler[[2]]$SingleR.single,
          monocytes$seurat@dr$tsne@cell.embeddings,do.label=F,
          do.letters =F, dot.size = 2)
out$p
```

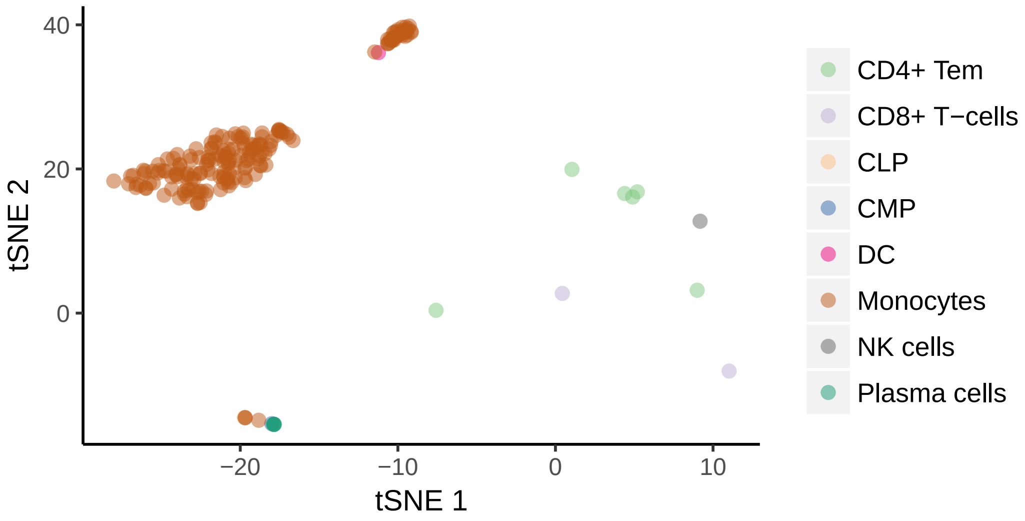

We can see that the t-SNE plot already suggests that there are 16 cells that are not part of the main cluster (which means that the sorting purity was ~92%). *SingleR* detected those cells to be plasma cells (4 cells), T-cells (8 cells), 2 progenitors, 1 DC and 1 NK cell. Is *SingleR* correct?

```
SingleR.DrawHeatmap(monocytes$singler[[2]]$SingleR.single,top.n = 20,
                    clusters=monocytes$singler[[2]]$SingleR.single$labels)
```

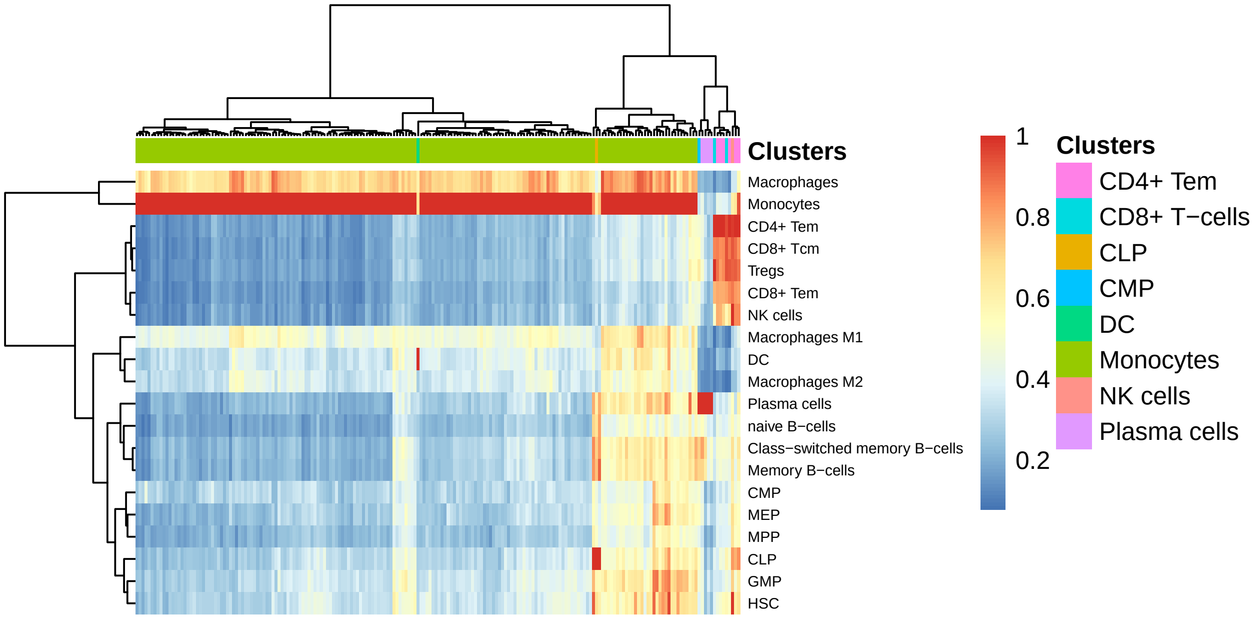

We can see that *SingleR* is quite convinced in its calls, giving low monocytes scores to those cells. Using markers for rare cell types (at least in the monocytes sorted cells) is problematic, since marker-based analysis is focused on clusters and not on single cells.

Interestingly, as in the t-SNE plot we see two distinct monocytes clusters, showing the ability to use *SingleR* for clustering.

**Example 4: GSE108097 (Mouse cell atlas) – Han et al. (2018)**

Using a method called Microwell-Seq, in this study the authors analyzed more than 400,000 single cells covering all of the major mouse organs. We reanalyzed the data from this data to allow visualization and exploration of the cellular heterogeneity of the different mouse organs. This effort allows the user a simple method to explore this invaluable comprehensive dataset. Here we present only one example from a bladder sample, and refer the user to the *SingleR* web tool for further evaluation of other organs.

```
load (file.path(path,'GSM2889480_Bladder.RData'))
out = SingleR.PlotTsne(singler$singler[[1]]$SingleR.single.main,
         singler$meta.data$xy,do.label=FALSE,
         do.letters =F,labels=singler$singler[[1]]$SingleR.single.main$labels,
         dot.size = 1, font.size = 8)
out$p
```

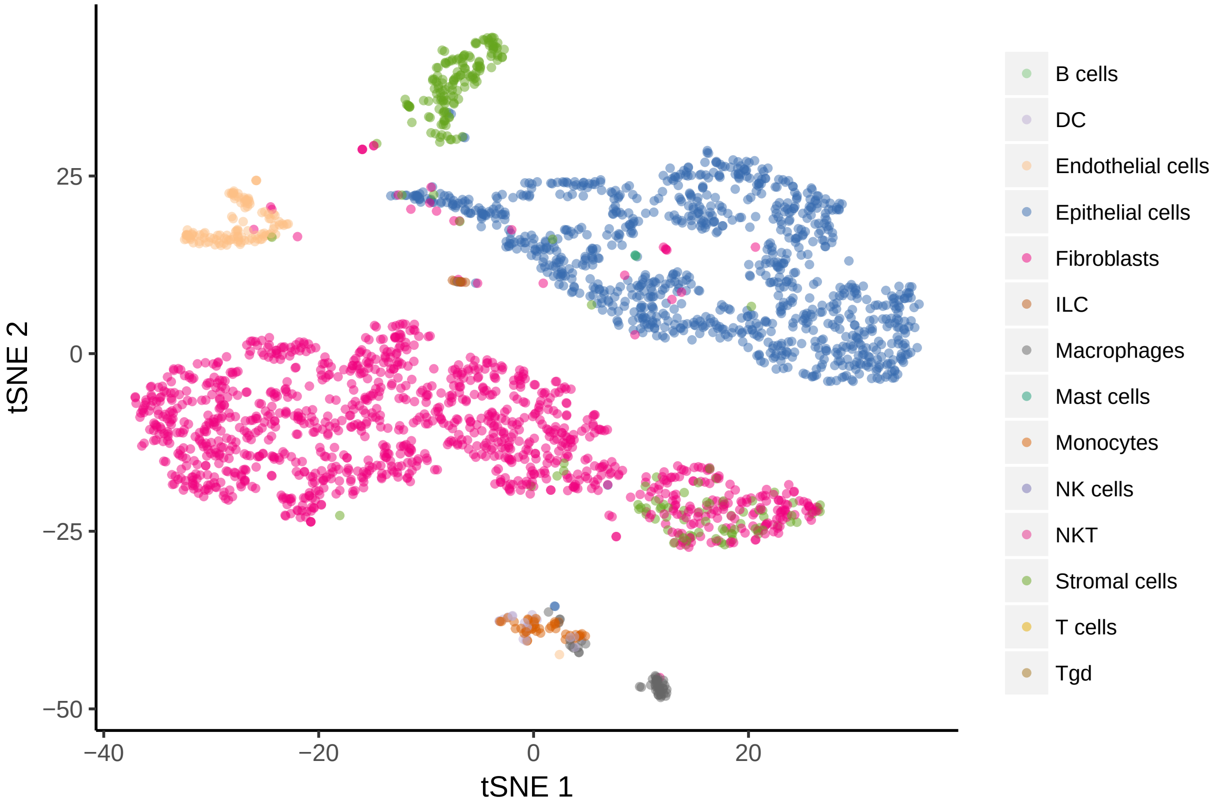

We can see that *SingleR* was able to resolve the cellular heterogeneity and allowing a rapid and unbiased approach to further explore the cellular heterogeneity in this organ.

#### ***SingleR*** **web tool**

The *SingleR* web tool contains >50 publicly available scRNA-seq datasets. All data has been reprocessed with the tools described above and the web tool allows the user immediate access to analyze the data and perform further investigations on published single cell data. In addition, we invite users to upload their own scRNA-seq data which will be analyzed on our servers and will process a SingleR object that can then be uploaded and used on the website (privately, only the user with the object is able to view it). Please visit http://comphealth.ucsf.edu/SingleR for more information.
